# Clinical-grade platelet-derived biomaterials mitigate mitochondrial dysfunction in preclinical models of neuronal injury

**DOI:** 10.1101/2025.09.11.675564

**Authors:** Kirti Gupta, Liling Delila, Ming-Li Chou, Nhi Thao Ngoc Le, Hsin-Yi Wu, David Blum, Anup Pandith, Thierry Burnouf

**Author notes:** **Corresponding Authors:** -Ming-Li Chou, Graduate Institute of Biomedical Materials and Tissue Engineering, College of Biomedical Engineering, Taipei Medical University, Taipei 110, Taiwan; -Thierry Burnouf, Graduate Institute of Biomedical Materials and Tissue Engineering, College of Biomedical Engineering, Taipei Medical University, Taipei 110, Taiwan.

## Abstract

Platelet-derived biomaterials are emerging as promising cell-free therapeutic platforms for regenerative medicine and neurorestoration. Among them, human platelet pellet lysates (HPPL) and platelet-derived extracellular vesicles (PEVs), prepared from clinical-grade allogeneic platelet concentrates, provide two complementary biomaterial formats: a soluble trophic factor-rich lysate and a vesicular formulation enriched in bioactive cargo. Because mitochondrial dysfunction and redox imbalance are hallmarks of neurodegeneration, we investigated whether these platelet-derived biomaterials could protect against rotenone-induced mitochondrial injury. HPPL and PEVs were bioprocessed from clinical-grade platelet concentrates and characterized for protein content, antioxidant capacity, vesicle morphology, size distribution, concentration and platelet and EVs markers. Differentiated N2A and SH-SY5Y neuronal cells were pretreated with 5% *v/v* HPPL or PEVs before rotenone (5 µM) exposure, while zebrafish embryos received 20 μg/mL HPPL or PEVs before rotenone (200 nM) challenge. Both biomaterials restored ATP production, reduced reactive oxygen species (ROS), preserved mitochondrial membrane potential and ultrastructure, and improved neuronal survival. They also normalized key markers of mitochondrial biogenesis and dynamics, including peroxisome proliferator-activated receptor gamma coactivator-1 alpha (PGC-1α), mitofusin-1 (MFN1), and dynamin-related protein 1 (DRP1). Proteomic analyses further showed enrichment of mitochondrial-associated antioxidant and metabolic proteins in HPPL and PEVs, and revealed restoration of oxidative phosphorylation, tricarboxylic acid cycle-related pathways, antioxidant defense, and mitochondrial dynamics in rotenone-injured cells following pretreatment. In zebrafish embryos, both biomaterials improved survival and hatching, reduced developmental abnormalities and oxidative stress, and preserved mitochondrial ultrastructure. These findings identify HPPL and PEVs as platelet-derived biomaterials with complementary mitochondrial protective activity, supporting their development as scalable cell-free biotherapeutic platforms for neurodegenerative disorders.

## 1. Introduction

Platelet-derived biomaterials are receiving increasing attention as cell-free therapeutic platforms for regenerative medicine and tissue repair beyond the classical hemostatic role of platelets [1–3]. Platelets are enriched in bioactive molecules, including antioxidants, neurotrophic growth factors, chemokines, cytokines, and extracellular vesicles, many of which can modulate inflammation, oxidative stress, cellular metabolism, and tissue remodeling [4, 5]. Platelet α-granules contain not only classical growth factors but also chemokines such as platelet factor-4 (PF4/CXCL4) and CCL5 (RANTES), which are increasingly implicated in neuroimmune communication and neuroprotection in the injured brain [6, 7]. Because platelet concentrates (PCs) are routinely collected, pathogen-screened, and manufactured by blood establishments worldwide, they provide a clinically relevant and scalable source material for the development of standardized platelet-derived biomaterials [1].

Among these biomaterials, human platelet pellet lysates (HPPL) and platelet-derived extracellular vesicles (PEVs) represent two complementary cell-free formats. HPPL is a soluble lysate enriched in trophic and redox-active factors released after controlled platelet disruption, whereas PEVs are membrane-enclosed nanoscale vesicles carrying proteins, lipids, and signaling molecules that may facilitate bioactive cargo delivery to recipient cells [8–11]. Both can be produced from clinical-grade PCs and contain neurotrophic factors, antioxidant enzymes, and other redox-active mediators [8–11]. These characteristics position HPPL and PEVs as promising secretome-derived biomaterials for neuroregenerative and neuroprotective applications [12, 13]. Previous studies from our group and others have shown that HPPL and platelet-derived EVs display neuroprotective, regenerative, and delivery-related activities in several preclinical models, supporting their value as translational platelet-derived biomaterials [9, 11, 14]. Importantly, HPPL and PEVs provide an opportunity to compare two complementary platelet-derived biomaterial formats generated from the same clinical source material: a soluble lysate enriched in freely available trophic and antioxidant factors, and a vesicular formulation in which bioactive cargo is presented within membrane-enclosed structures [8, 9, 15]. Previous observations further suggest that these two formats may exert overlapping but distinct biological effects, making their direct comparison particularly relevant for biomaterial development [16].

Their translational potential is further supported by the broad accessibility of clinical-grade PCs, which could enable development of locally sourced biotherapies even in regions with limited capacity for advanced manufacturing [17]. Plans for upcoming clinical trials in amyotrophic lateral sclerosis (ALS), a disorder strongly linked to mitochondrial dysfunction, further highlight the therapeutic interest of platelet-derived biotherapies [14].

This biomaterial concept is particularly relevant in neurodegenerative disorders, where mitochondrial dysfunction and oxidative stress are central pathogenic mechanisms. Mitochondria regulate ATP production through oxidative phosphorylation (OXPHOS), as well as reactive oxygen species (ROS) homeostasis, calcium signaling, apoptosis, and quality-control pathways such as biogenesis, fission, fusion, and mitophagy [18] [19]. In neurons, which are highly dependent on oxidative metabolism, disruption of these processes leads to bioenergetic failure, ROS accumulation, defective mitochondrial dynamics, and ultimately cell death [20] [21] [22, 23] [24, 25]. Such alterations are common to several central nervous system (CNS) disorders, including Alzheimer’s disease, Parkinson’s disease, amyotrophic lateral sclerosis, and Huntington’s disease [26, 27]. Therefore, biomaterials capable of delivering antioxidant and mitochondrial-supportive cargo may provide a promising cell-free strategy to restore neuronal resilience under pathological stress.

Rotenone, a mitochondrial complex I inhibitor, is widely used to model mitochondrial dysfunction because it reproduces key features of oxidative phosphorylation impairment, ROS overproduction, mitochondrial depolarization, and ultrastructural damage *in vitro* and *in vivo* [28] [29] [30]. In this context, HPPL and PEVs offer an attractive therapeutic rationale. HPPL may provide a concentrated soluble pool of trophic and antioxidant factors, whereas PEVs may additionally enable vesicle-mediated uptake and intracellular trafficking of bioactive cargo [8–11]. Their shared origin from clinical-grade PCs further supports their value as scalable platelet-derived biomaterials for translational use [1].

In the present study, we investigated whether HPPL[15] and PEVs [9] can mitigate rotenone-induced complex I inhibition, oxidative stress, mitochondrial dysfunction, and ultrastructural damage in neuronal cell models and zebrafish embryos. By combining physicochemical characterization, functional mitochondrial assays, imaging, and proteomic analyses, we compared how these two platelet-derived biomaterial formats influence bioenergetics, redox balance, mitochondrial dynamics, and structural preservation. Our findings provide additional mechanistic insight into the therapeutic potential of platelet-derived biomaterials and support their development as scalable cell-free interventions for neurodegenerative disorders.

## 1. Materials and Methods

### 2.1. Source of Platelet Concentrates (PCs)

Clinical-grade human PCs were collected from healthy voluntary donors using Haemonetics MCS^®^+ multicomponent apheresis systems, following standard protocols at the Taipei Blood Center, Taiwan Blood Service Foundation (Taipei, Taiwan), with approval from the Institutional Review Board of Taipei Medical University (TMU-JIRB N201802052). PCs were anticoagulated with a citrate-based solution and suspended in 100% plasma. Upon reaching their transfusion shelf-life (5 days post-collection), the PCs were transported to the Taipei Medical University (TMU) laboratory under controlled temperature conditions and maintained under gentle agitation at 22°C ± 2 °C. Within 24 hr of receipt, the PCs were centrifuged at 300 x g for 15 min to sediment and remove any residual RBCs. Blood cell count was determined using the ABC Vet blood cell counting system (ABC Diagnostics, Montpellier, France).

### 2.2. Bioprocessing of heat-treated platelet pellet lysates (HPPL) and platelet derived extracellular vesicles (PEVs)

HPPL and PEVs were prepared by pooling five batches of PCs as previously reported [9, 15]. Briefly, PCs were first centrifuged at 3,000 x g for 30 min at 22 ± 2°C to pelletize the platelets. Pellets were gently washed and resuspended in 0.1 µm filtered phosphate-buffered saline (PBS) into a volume equal to 10% of the original PCs volume to prepare HPPL. The respective supernatant containing plasma was carefully recovered for the preparation of PEVs. The resuspended pellets were lysed by three cycles of freeze (-80 ± 1°C) and thaw (37 ± 1°C), followed by centrifugation at 4,500 x g for 30 min to remove any remaining intact platelets and cell debris. The resulting platelet pellet lysate (PPL) was heat-treated at 56 ± 1°C for 30 min to inactivate or remove pro-coagulant factors [8] and fibrinogen [15] and immediately cooled in an ice bath for 5 min and clarified by centrifugation at 10,000 x g for 15 min to obtain HPPL. PEVs were isolated from the previously recovered PCs supernatant by centrifugation at 6,000 × g for 30 min at 25 ± 2°C to remove the cell debris and ultracentrifugation (Beckman L-90K Ultracentrifuge) of the supernatant at 30,000 × g for 3 hr at 18 ± 2°C to pelletize the PEVs. The PEVs were washed and subsequently resuspended in a volume of PBS equal to 1% of the initial PCs volume. The HPPL and PEVs preparations were stored at -80°C. They were thawed at 37°C and centrifuged at 6000 × g for 15 min at 4°C to remove any precipitates before use.

### 2.3. Physicochemical Characterization of HPPL and PEVs

The total protein concentration of HPPL and PEVs was determined using the bicinchoninic acid (BCA) assay (Thermo Scientific, #23225), following the manufacturer’s instructions. Samples were diluted 10-fold, mixed with BCA working reagent, and incubated at 37°C for 30 min. Absorbance was measured at 562 nm using a microplate reader (BioTek, EPOCH2). For PEV-specific characterization, multiple analytical approaches were employed. The morphology of PEVs was visualized using cryo-electron microscopy (cryo-EM). Dynamic Light Scattering (DLS) was used to assess the hydrodynamic size distribution of PEVs and measurements were performed using a Zetasizer Nano ZS (Malvern Instruments, UK) in 100 µL disposable cuvettes, as previously described [26, 30]. Nanoparticle Tracking Analysis (NTA) was carried out using the NanoSight NS300 system (Malvern Instruments, UK) to measure particle size distribution and concentration. PEVs were diluted 10,000-fold in PBS, and 800 µL of the sample was injected via syringe pump. The particles were illuminated with a laser and recorded for 60 sec to determine size and count based on Brownian motion. Samples were applied to glow-discharged holey carbon grids using an FEI Vitrobot, blotted, and vitrified in liquid nitrogen. Imaging was performed at 50,000-fold magnification using an FEI Tecnai F20 microscope operated at 200 kV at the cryo-EM facility of Academia Sinica, Taiwan. The Exo-Check Exosome Antibody Array (EXORAY210B-8, System Biosciences) was used to confirm the presence of characteristic exosomal surface markers. A total of 50 µg of PEVs protein was lysed, labeled with detection reagent, and incubated overnight at 4°C on the antibody membrane. After washing, membranes were treated with an HRP-conjugated detection mix for 30 min, developed using Clarity Max Western enhanced chemiluminescence (ECL) substrate, and imaged using the Invitrogen iBright CL750 system. Band intensities were quantified after subtracting the background signal. The protein expression levels of PEVs were assessed using SDS-PAGE and Coomassie Brilliant Blue (ab119211, Abcam) staining was performed to observe the overall protein profiles of PEVs. For western blot analysis, proteins were transferred onto polyvinylidene difluoride (PVDF) membranes and membranes were blocked with 5% (w/v) non-fat milk in Tris-buffered saline containing 0.1% Tween-20 (TBST) for 1 hr at RT, followed by overnight incubation at 4°C with primary antibodies against CD42a (ab133573, Abcam), CD41 (ab134131, Abcam), and CD62P (ab255822, Abcam) at 1:1000 dilution each and β-actin used as control (GTX109639, Gentex). After washing with TBST, membranes were incubated with horseradish peroxidase (HRP)-conjugated anti-rabbit secondary antibodies for 1 hr at RT. Protein bands were visualized using an ECL detection kit and imaged with a ChemiDoc Imaging System.

### 2.4. Assessment of protective effects of HPPL and PEVs in N2A and SH-SY5Y neuronal cells

#### 2.4.1. Cell culture maintenance, Differentiation and Treatment protocol

The human neuroblastoma SH-SY5Y cells were kindly provided by David Devos (University of Lille, France), and mouse Neuro-2A (N2A) cells were obtained from the Bioresource Collection and Research Center (BCRC), Taiwan. SH-SY5Y and N2A cells were cultivated in Minimum Essential Medium/F-12 (MEM/F12; PM151220, Gibco) and high glucose Dulbecco’s modified eagle medium (DMEM, SH30243.02, Hyclone) respectively, both supplemented with 10% FBS (A4736301, Gibco) and 1% Antibiotic-Antimycotic (30-004-CI, Corning) in Nunc Cell Culture Flasks™ (156499, Thermo Scientific) incubated at 37°C and humidified with 5% CO2 and 95% air. Upon reaching 80-90% confluency, both cells were passaged every 3-4 days and seeded at 1 × 10⁶ cells. All experiments were performed using differentiated neurons (15-20 passaged cells) using 10 μM retinoic acid (RA) in the respective serum-deprived medium for 3-4 days. Rotenone (*83-7-94, ≥ purity 99.9%, Sigma*) was initially diluted in dimethyl sulfoxide (DMSO, Sigma-Aldrich) to a concentration of 1 mM (*v/v*) and further diluted in culture medium to final concentrations of 1 µM, 2 µM, 5 µM, and 10 µM to identify the lowest concentration that significantly reduced cell viability in 24 hr. Stock solutions of MitoQuinone Mesylate (MitoQ) (HY-100116A, MedChemExpress) were prepared in DMSO with the final concentrations in the culture medium being ≤ 0.1% (*v/v*). Based on preliminary dose-response findings, the experimental groups were further categorized in both SH-SY5Y and N2A cells as follows: Untreated cells (Control), cells treated with 5 µM rotenone for 24 hr to induce mitochondrial damage (Rot), cells pre-treated 1 hr with 100 nM MitoQ prior to 24 hr rotenone (MitoQ + Rot), cells pre-treated with 5% HPPL (*v/v*) for 1 hr prior to rotenone exposure (HPPL + Rot), cells pre-treated with 5% PEVs *(v/v)* for 1 hr prior to rotenone exposure (PEVs + Rot).

#### 2.4.2. Cellular uptake of HPPL and PEVs by cells

To assess cellular uptake of HPPL proteins and PEVs, both were fluorescently labeled using the Alexa Fluor™ 488 Protein Labeling Kit (A-10235, Thermo Fisher Scientific) at a final protein concentration of 2 mg/mL. N2A cells were seeded on sterilized cover slips, and after 24 hr, incubated with Alexa Fluor 488-labeled HPPL and PEVs in serum-free media for 1 hr to evaluate uptake. Following incubation, cells were washed thrice with pre-warmed PBS to remove unbound material, and subsequently stained with 100 nM MitoTracker™ Red CMXRos (M7512, Thermo Fisher Scientific) for 45 min to visualize mitochondria. Cells were then fixed with 4% paraformaldehyde (PFA), permeabilized with 0.5% Triton X-100, and counterstained with DAPI (4′,6-diamidino-2-phenylindole) (D9542, Sigma) for 10 min to visualize nuclei. Confocal imaging was performed using Stellaris 8 with 3D acquisition settings, and fluorescence intensities were quantified using ImageJ software. For colocalization analysis, the Coloc 2 plugin in ImageJ was used to calculate Pearson’s correlation coefficient, providing a quantitative measure of the spatial overlap between HPPL/PEVs signals and the mitochondrial network. The coefficient ranges from -1 to +1, where values closer to +1 indicate a strong positive correlation (high colocalization), values around 0 suggest no correlation (random distribution), and values near -1 indicate an inverse correlation.

#### 2.4.3. Cell viability assays

For assessing cell viability and distinguishing live and dead cells, two complementary assays were used. The colorimetric cell counting kit-8 (CCK-8 assay, 96992, Sigma-Aldrich), where 100 µL of cell culture medium received 10 µL of CCK-8 solution, which produces formazan in proportion to the viable cell count. After 4 hr incubation at 37°C, the absorbance at 450 nm was measured using a microplate reader (BioTek, EPOCH2). Viability was calculated as a percentage relative to untreated control cells using the formula: [O.D. of sample − O.D. of blank/O.D. of control − O.D. of blank] × 100. The live/dead double staining kit was utilized for simultaneous staining of Calcein-AM stained viable cells (green fluorescence), while propidium iodide (PI) stained dead cells (red fluorescence). After preparing the assay solution (10 µL solution A + 5 µL solution B in 5 mL PBS), cells were washed with PBS to remove any residual esterase activity, followed by incubation with the assay solution for 15 min at 37°C. Advance fluorescence microscope (Leica DMi8) was used to simultaneously monitor live cells at 490 nm excitation and visualize dead cells at 545 nm excitation.

#### 2.4.4. Complex I Enzyme Activity Assay

Mitochondrial Complex I activity was quantified using the Complex I Enzyme Activity Assay Kit (ab109721, Abcam) as per the manufacturer’s instructions. Briefly, N2A cells from each experimental group were washed twice with cold PBS and harvested by centrifugation at 500 × g for 5 min at 4°C. The cell pellets were lysed with kit-supplied 10% detergent for 30 min on ice and protein concentrations were determined. Lysates were centrifuged at 16,000 × g for 20 min at 4°C, and supernatants were collected. For each condition, 125 μg of protein was loaded per well to a final volume of 200 µL with incubation Solution, in a 96-well plate, and incubated for 3 hr at RT to allow Complex I capture. Following three washes, 200 µL of assay solution was added to each well and immediately, the absorbance was recorded at 450 nm for 30 min at 1 min intervals with intermittent shaking using a microplate reader (BioTek, EPOCH2). Data were normalized to total protein concentrations and expressed relative to control levels.

#### 2.4.5. Cellular bioenergetic assays

To reflect the cellular metabolic state in N2A cells, the ATP production was assessed using the ATP detection assay kit (ab83355, Abcam) and lactic acid concentration levels were determined using the lactic acid assay kit (ab65331, Abcam). Briefly, 1 x 10^6^ N2A cells were harvested, washed in cold PBS and resuspended in 100 μL of ATP assay buffer or Lactate Assay Buffer, respectively. Cells were homogenized and centrifuged at 13,000 × g at 4°C for 2-5 min. Supernatant containing the cellular contents were collected on ice and deproteinized using the PCA/KOH protocol following the kit instructions. After cell sample preparation, 50 µL of the supernatant from each sample and freshly prepared ATP or lactate standards were mixed with ATP or lactate reaction mix and loaded individually into a 96-well plate in triplicate. The plates were incubated at RT for 30 min, protected from light, and absorbance for ATP was measured at 570 nm and lactate was measured at 450 nm using a microplate reader (BioTek, EPOCH2). The intracellular ATP and lactate concentrations were determined from three biological replicates based on the mean optical density using the respective standard curves.

#### 2.4.6. Real time oxygen consumption rate (OCR) quantification

The direct measurement of the oxygen consumption rate (OCR) in live N2A cells was performed using XFe24 Extracellular Flux Analyzer-2 (Seahorse Bioscience, Taiwan), modifying the essential manufacturer’s instructions from Seahorse XF Cell Mito Stress Test kit (103015-100, Agilent). Cells were seeded in specialized Seahorse XF Cell Culture microplates at an optimal density of approximately 1.5 × 10⁴/well in 100 μL culture media for 2-3 hr to allow cell attachment. The medium was then adjusted to 250 µL per well and incubated for 24 hr. To avoid interference from the kit-supplied rotenone, oxidative stress was induced using hydrogen peroxide (H₂O₂) at a previously standardized dose of 100 µM. Therefore, the following experimental groups were established: untreated cells, cells treated with 100 µM H_2_O_2_ for 24 hr, cells pre-treated with 100 nM MitoQ for 1 hr, followed by 100 µM H_2_O_2_ for 24 hr, cells pre-treated with 5% (v/v) HPPL for 1 hr, followed by 100 µM H_2_O_2_ for 24 hr, cells pre-treated with 5% (v/v) PEVs for 1 hr followed by 100 µM H_2_O_2_ for 24 hr. One hour before measurement, culture medium was replaced with Seahorse XF assay medium and sequential injections of modulators were used during the assay to evaluate different components of mitochondrial respiration: Oligomycin (1 µM) inhibited ATP synthase to assess ATP-linked respiration; FCCP (0.5 µM), a mitochondrial uncoupler, was used to determine maximal respiration; and Antimycin A/Rotenone (0.5 µM) inhibited complexes III and I to measure non-mitochondrial respiration. OCR was recorded in real time before and after each injection to quantify basal respiration, ATP production, maximal respiration, and spare respiratory capacity. Data were exported using the Wave desktop software to generate XF-Cell Mito Stress Test reports ad normalized to protein concentrations.

#### 2.4.7. Live cell imaging to assess cellular and mitochondrial reactive oxygen species (ROS)

Intracellular ROS levels in N2A and SH-SY5Y cells were measured using a 2′,7′-dichlorodihydrofluorescein diacetate (DCFDA) cellular ROS Assay Kit (ab113851, Abcam). Each experimental group of cells seeded in a 35 mm dish was incubated with 10 μM DCFDA for 45 min at 37°C, followed by three washes with kit-supplied 1X buffer. The oxidative conversion of DCFDA into its fluorescent form was analyzed to determine the ROS levels. For detecting mitochondria-specific ROS levels, MitoSOX Red superoxide indicator (HY-D1055, MedChemEexpress) was used to selectively target mitochondrial superoxide in living cells. The initial 1 mM stock was prepared in DMSO and diluted to a final working concentration of 5 μM in HBSS. Cells were incubated for 30 min at 37°C, followed by three washings to remove unbound dye. Fluorescence signal intensity was assessed under advanced fluorescence microscopy (LEICA DMi8) within 2 hr and quantified using ImageJ software.

#### 2.4.8. Determination of Glutathione peroxidase (GPx)

To measure the NADPH consumption in the enzyme-coupled reactions, SH-SY5Y cells (2 x 10^6^) were harvested, washed in cold PBS and lysed with 200 μL of ice-cold assay buffer. Cells were homogenized and centrifuged at 10,000 × g at 4°C for 15 min and the supernatant was collected for analysis using the Glutathione Peroxidase Assay Kit (MAK437, Merck). In 96-well plate, 40 µL of reaction mix was added to each sample well (100 µL), positive control (50 µL) and reagent control wells (50 µL) and incubated at 25°C for 15 min. Afterwards, cumene hydroperoxide solution (10 µL) was added to the reaction wells and A1 was measured at T1 on a microplate reader (BioTek, EPOCH2) at OD340 nm, followed by A2 measured at T2 after 5 min. All experiments were performed in duplicate, with three independent experiments, and calculations were carried out according to the kit guidelines.

#### 2.4.9. Oxygen radical absorbance capacity (ORAC) of HPPL and PEVs

The antioxidant capacity of HPPL and PEVs was evaluated using the oxygen radical absorbance capacity (ORAC) assay (Zen-Bio, AOX-2). In a 96-well plate format, 150 µL of fluorescein working solution was added per well, followed by 25 µL of Trolox standards, 100-fold diluted HPPL and PEVs samples, or positive controls. The positive control included human plasma (60-65 mg/mL, used at 100-fold dilution), ATP (ab83355, Abcam) and MitoQ (MitoQ10, HY-100116A, MedChemExpress), both used at 10 μM final concentration in duplicates. After a 15-min incubation at 37°C, the reaction was initiated by adding 25 µL of AAPH solution. Fluorescence (Ex/Em: 485/530 nm) was recorded at one-min intervals for 30 min using a Thermo Varioskan Flash spectrofluorometer. The area under the curve was calculated, and antioxidant capacity was expressed as Trolox equivalents.

#### 2.4.10. Functional and Ultrastructural assessment of Mitochondria

To determine the changes in mitochondrial membrane potential (ΔΨm) of cells, the JC-1 Mitochondrial Membrane Potential Assay Kit (HY-K0601, Med Chem Express) was used according to the vendor’s protocol. JC-1 (5,5’, 6,6’-Tetrachloro-1,1’, 3,3’-tetraethyl-amidinephthalocyanine iodide) exists as monomers when the membrane potential is low (△Ψm↓), emitting green fluorescence, whereas it generates J-aggregates, when the membrane potential is high (△Ψm↑), showing red fluorescence. N2A cells from each treatment group were stained with 2 μM JC-1 probe for 15-20 min at 37℃ in an incubator and washed with 1X PBS thrice. The intensities of red and green fluorescence were observed using fluorescence microscopy and quantified for mitochondrial depolarization using ImageJ software. Additionally, these N2A cells from each treatment group were also incubated with 100 nM MitoTracker Red for 30 min at 37 °C to visualize mitochondria based on membrane potential, viewed under a ZEISS LSM900 confocal microscope. Transmission electron microscopy (TEM; HT-7700, HITACHI) was used to examine the subcellular mitochondrial ultrastructure in each experimental set of SH-SY5Y cells. Briefly, 5 × 10⁴ cells were seeded per group. After 24 hr of treatment, cells were fixed in a solution containing 2% PFA and 2.5% glutaraldehyde in 0.1 M cacodylate buffer for 1 hr at RT. Post-fixation was performed with 1% osmium tetroxide (OsO₄) for 1 hr, followed by three washes in 0.1 M cacodylate buffer supplemented with 7% sucrose. Samples were then dehydrated through a graded ethanol series (70%, 80%, 90%, and 100%), each step for 15 min. Cells were embedded in epoxy resin, and ultrathin sections (∼50 nm) were obtained using an ultramicrotome. Sections were subsequently stained with uranyl acetate before imaging under the TEM.

#### 2.4.11. PGC-1α quantification by ELISA

To evaluate mitochondrial biogenesis, peroxisome proliferator-activated receptor gamma coactivator-1 alpha (PGC-1α) levels were measured using a mouse PGC-1α ELISA kit (Sunlong, SL0797Mo) following the manufacturer’s protocol. After treatment, N2A cells were washed with cold PBS, trypsinized, and centrifuged at 2,000 rpm for 5 min at 4°C. Pellets were washed twice with cold PBS and resuspended to a final density of 1 × 10⁶ cells/mL in PBS. Cells were subjected to three freeze–thaw cycles (–80°C/37°C). Lysates (clear supernatant) were collected for ELISA after centrifugation at 2,500 rpm for 20 min. 50 uL of sample and standards were added to a 96-well plate pre-coated with anti-PGC-1α antibodies and incubated for 90 min at 37°C. After washing, the HRP-conjugate reagent was added and incubated at 37°C for 30 min. Color was developed by adding 50 µL each of Chromogen Solutions A and B, incubating in the dark at 37°C for 15 min, and finally, the reaction was terminated with 50 µL stop solution. Absorbance was measured at 450 nm using a microplate reader (BioTek, EPOCH2). PGC-1α levels were calculated from a standard curve and normalized to protein concentration determined by BCA assay.

#### 2.4.12. Immunofluorescence

N2A cells were fixed with 4% PFA for 20 min at 4°C, followed by permeabilization with 0.1% Triton X-100 for 10 min at RT. Cells were blocked with 2% bovine serum albumin (BSA) for 1 hr at RT. Immunostaining was performed by incubating the cells overnight at 4°C with primary antibodies targeting PGC-1α (1:50, sc-518038, Santa Cruz), Mitofusin-1 (1:100, ab126575, Abcam), and TOMM20 (1:100, ab186735, Abcam). After three PBS washes, cells were incubated with Alexa Fluor 488- or 594-conjugated anti-rabbit or anti-mouse secondary antibodies (1:1000, Invitrogen) for 1 hr at RT. Finally, samples were mounted using DAPI-containing mounting medium, and fluorescence images were acquired using a ZEISS LSM900 confocal microscope.

#### 2.4.13. Sample Preparation for Cellular Proteomic Analysis

N2A cell samples from control, rotenone treated, HPPL + rotenone, and PEVs + rotenone groups were processed for proteomic analysis. Cells were harvested, washed twice with ice-cold PBS, and lysed to extract total cellular proteins. Protein concentration was quantified using the BCA assay, and samples were normalized to a final concentration of ≥1 μg/μL. Protein precipitation was performed using cold acetone (−20 °C, pre-cooled ≥24 h) at a 1:4 (v/v) ratio with cell lysates. Samples were vortexed briefly, incubated overnight at −20 °C for protein precipitation, and subsequently submitted to the Proteomics Core Facility at National Taiwan University (NTU) for liquid chromatography–tandem mass spectrometry (LC–MS/MS) analysis.

### 2.5. Assessments in Zebrafish (ZF) embryos

#### 2.5.1. Experimental ZF maintenance, Embryos harvesting and Drug exposures

Adult wild-type ZF (AB strain, 8–12 months old) were obtained from TMU’s ZF Core Facility and maintained in a 90 × 60 × 45 cm circulating tank at 28°C under a 14:10 hr light–dark cycle, with pellet feeding twice daily. All husbandry and experimental procedures adhered to the Taiwan Animal Protection Act. As per regulatory guidelines, no approval was required for experiments involving embryos up to 96 hours post-fertilization (hpf). Fertilized embryos were collected daily from 9:00 to 10:00 am after natural spawning and incubated in artificial freshwater (0.2 mM CaSO₄, 0.5 mM NaCl, 0.2 mM MgSO₄, 0.16 mM KH₂PO₄, 0.16 mM K₂HPO₄; pH 6.8–7.0). At 4 hpf, embryos with normally developed blastula were randomly distributed in 24-well plates (10 embryos/well) containing 1 mL of E3 medium and incubated at 28.5°C. Zebrafish Embryo Toxicity (ZFET) assay was performed using diluted stocks of HPPL and PEVs based on protein concentration, and rotenone at concentrations from 50 nM to 1000 nM. Based on survival analysis and prior literature [4], treatment groups were selected as follows (n = 30 per group) - Group I: Control; Group II: 200 nM rotenone (Rot); Group III: 20 μg/mL HPPL pre-treatment 1 hr before Rot (HPPL + Rot); Group IV: 20 μg/mL PEVs pre-treatment 1 hr before Rot (PEVs + Rot). Fresh media and treatments were replaced every 24 h for five days.

#### 2.5.2. Determination of development index, hatch time and morphological assessment

To record survival curves, each treatment group was observed at 4, 24, 48, 72, 96 hpf and dead embryos were removed from the plates immediately. For the calculation of hatching rate, 48 and 72 hpf embryos hatched into larvae were assessed in relation to the total egg count. The cumulative mortality and developmental malformations of embryos and larvae were observed every 24 hr. To observe morphological abnormalities, embryos from each group were anesthetized with MS-222 for 2-3 min until immobile and bright-field images were captured using a fluorescence stereomicroscope (ZEISS) at 4, 24, 48, 72, 96 hpf. The embryonic developmental parameters for malformation rate were monitored for pericardial edema (PE), which is swelling around the heart, yolk sac edema (YE), swelling of the yolk sac, axis malformation (AM), abnormal body axis development, and bent tail (BT), curvature of the tail.

#### 2.5.3. Biodistribution of HPPL and PEVs in whole embryo and mitochondrial uptake

ZF embryos were exposed to selected doses of AlexaFluor 488-labeled HPPL and PEVs in E3 medium for 1 hr, protected from light. After treatment, embryos were washed with E3 medium three times and stained with 500 nm MitoTracker Red for 1 hr to visualize mitochondria-rich cells. The embryos were stained with 1-5 µg/mL DAPI for 30min to visualize nuclei, and imaged using laser Stellaris 8 confocal microscopy at 60-fold magnification. Imaging was performed at 24 hpf and 96 hpf to monitor the distribution and uptake of HPPL and PEVs and their localization within mitochondria. The fluorescence intensities were measured using ImageJ software.

#### 2.5.4. Oxidative stress determination, mitochondrial isolation and functional assays

Intracellular ROS production of ZF embryos was assessed using DCFH-DA. Live ZF embryos from each treatment group at 96 hpf were incubated with 10 μM of DCFH-DA for 30 min at RT and observed using a confocal microscope (ZEISS LSM900). A pool of 30 ZF embryos per treatment group was collected and mitochondria were isolated using the Mitochondrial Isolation Kit (ab288084, Abcam) following the manufacturer’s protocol. Briefly, embryos were washed twice and homogenized in mitochondrial extraction buffer supplemented with protease inhibitor. The homogenate was centrifuged at 600 x g for 10 min at 4°C to remove debris, and the supernatant was centrifuged at 7,000 x g for 10 min at 4°C to pellet the mitochondria. Mitochondria were washed and collected by centrifugation and resuspended in Storage Buffer. The protein concentration was determined and isolated mitochondria were stored at -80°C until use. The ATP content of isolated mitochondria compared to homogenized embryos was measured on a microplate reader (BioTek, EPOCH2) using the ATP detection assay kit (ab83355, Abcam).

#### 2.5.5. Transmission Electron Microscopy

ZF Embryos at 96 hpf from each treatment group were fixed with 2.5% glutaraldehyde in 0.1 M PBS (pH 7.4) overnight at 4°C. The fixed embryos were then rinsed three times in PBS and subsequently post-fixed with 1% osmium acid for 1 hr. After three times washing in distilled water, stained in 2% uranyl acetate for 30 min, and then dehydrated in a graded series of ethanol (50% for 15 min, 70% for 15 min, 90% for 15 min, 100% for 20 min) and 100% acetone for 20 min twice. The embryos were treated in Spurr’s resin and acetone (1:1) for 2 hr at RT and then embedded in Spurr’s resin at the opposite position. Ultra-thin sections (50 nm) through the embryonic heads were obtained using a glass knife on a Leica Reichert microtome and placed on formvar-coated copper grids. Sections were stained with uranyl acetate and viewed under TEM. Mitochondrial area (μm^2^) was quantified and the circulatory index (0-1) was used to estimate the swelling index using ImageJ. Cristae morphology was scored from 0 to 4 based on ultrastructural features as shown in table 1 [31]:

**Table 1.** Scoring system for mitochondrial cristae morphology.

|  |  |
| --- | --- |
| Score 0 | Absent or no sharply defined cristae |
| Score 1 | Very few cristae (>10%) |
| Score 2 | Few cristae, disorganized (>25%) |
| Score 3 | Moderate cristae, partially aligned (>50%) |
| Score 4 | Dense, well-organized regular cristae |

### 2.6. Statistical analysis

All the raw data were compiled in Excel and Google Sheets. Data are expressed as the mean ± standard deviation (SD) where n = 6 biological independent replicates unless stated otherwise. Comparisons were performed using the Student’s *t* test, paired *t* test comparing two conditions, with one-way ANOVA comparisons tests (Tukey’s, or Dunnett’s test). Survival analysis was performed using Kaplan-Meier Curve, presented using Prism software (GraphPad Prism, version 8.02).

## 3. Results

### 3.1. Bioprocessing and biophysical characterization of HPPL and PEVs

PCs had platelet counts ranging from 973–1418 (× 10^3^/μL), corresponding to a total of 2.7–3.5 × 10^11^ platelets. The residual red blood cells (RBCs) and white blood cells (WBCs) concentrations were both below detectable limits (< 0.7 × 10^6^/μL and < 0.7 × 10^6^/μL, respectively), ensuring minimal contamination from other blood cell types. HPPL and PEVs were isolated through a differential centrifugation protocol, as outlined in Figure 1, by pooling preparations obtained from five separate batches of clinical-grade PCs to ensure consistency. Total protein content measured using the BCA assay, averaged 7.53 mg/mL for HPPL and 2.36 mg/mL for PEVs (Figure 2A). Cryo-EM revealed that PEVs were membrane-bound vesicles ranging from 50 to 300 nm in diameter, consistent with the typical vesicular morphology (Figure 2B). DLS analysis showed that PEVs exhibited a dominant population of nearly 200 nm, along with a smaller population around 20 nm (Figure 2C). NTA further confirmed the presence of vesicles in the range of 50 – 250 nm, with particle concentrations between 10^11^ – 10^12^ particles/mL in PEVs (Figure 2D). Marker profiling using a human-specific exosome antibody array confirmed the presence of classical extracellular vesicle (EV) markers in PEVs such as tumor susceptibility gene 101 (TSG101), programmed cell death 6 interacting protein (ALIX), tetraspanin (CD63 and CD81), Annexin A5 (ANXA5), flotillin-1 (FLOT1) and epithelial cell and intracellular adhesion proteins (EpCAM, ICAM) along with the negligible cellular contamination assessed by Golgi marker (GM130) (Figure 2E & F). SDS-PAGE followed by Coomassie brilliant blue staining revealed the protein expression patterns of isolated PEVs (Figure 2G). Western blot analysis further confirmed the presence of platelet-specific glycoproteins CD42a and CD41, as well as the platelet activation marker CD62P, in PEVs lysates (Figure 2H).

**Figure 1.**
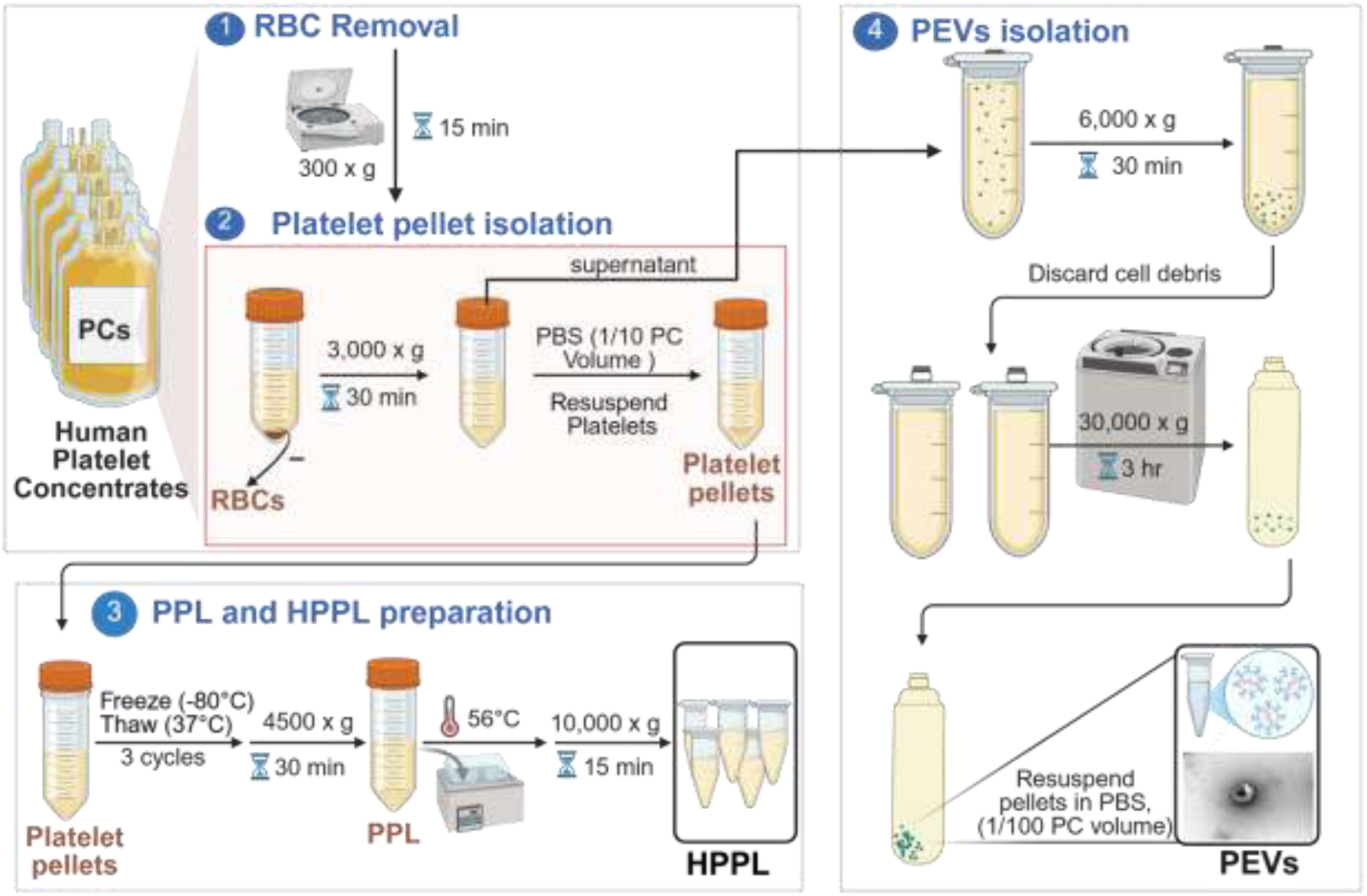
Schematic representation of HPPL and PEVs isolation from PCs. (1) PCs were centrifuged to remove residual RBC contaminants, and the platelet count was measured. (2) PCs were centrifuged to pelletize the platelets and resuspend them in PBS. (3) The suspended platelet pellet underwent three freeze-thaw cycles followed by centrifugation to obtain PPL that was heat-treated, then cooled down and centrifuged to generate HPPL. (4) PEVs were isolated from the corresponding supernatant by differential centrifugation and ultracentrifugation, resuspended in PBS. The TEM image illustrates the desired spherical PEVs obtained following the described protocol. Note: HPPL and PEVs were aliquoted and stored at -80*°*C until use. Abbreviations: PC: Platelet concentrate, RBCs: Red blood cells, PBS: Phosphate buffer saline, PPL: Platelet pellet lysate, HPPL: Human platelet pellet lysate, PEVs: Platelet derived extracellular vesicles. *Created in* https://BioRender.com.

**Figure 2.**
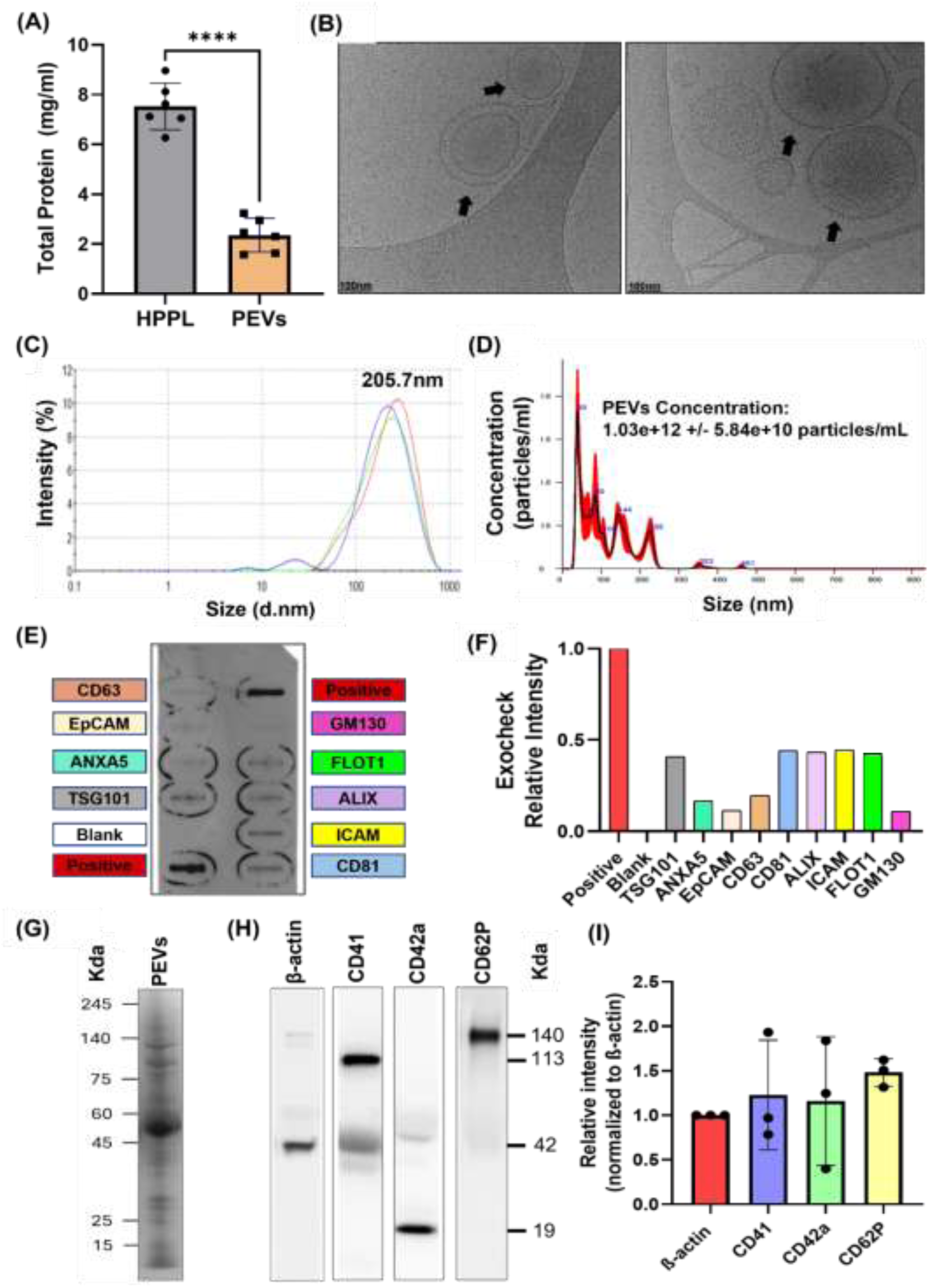
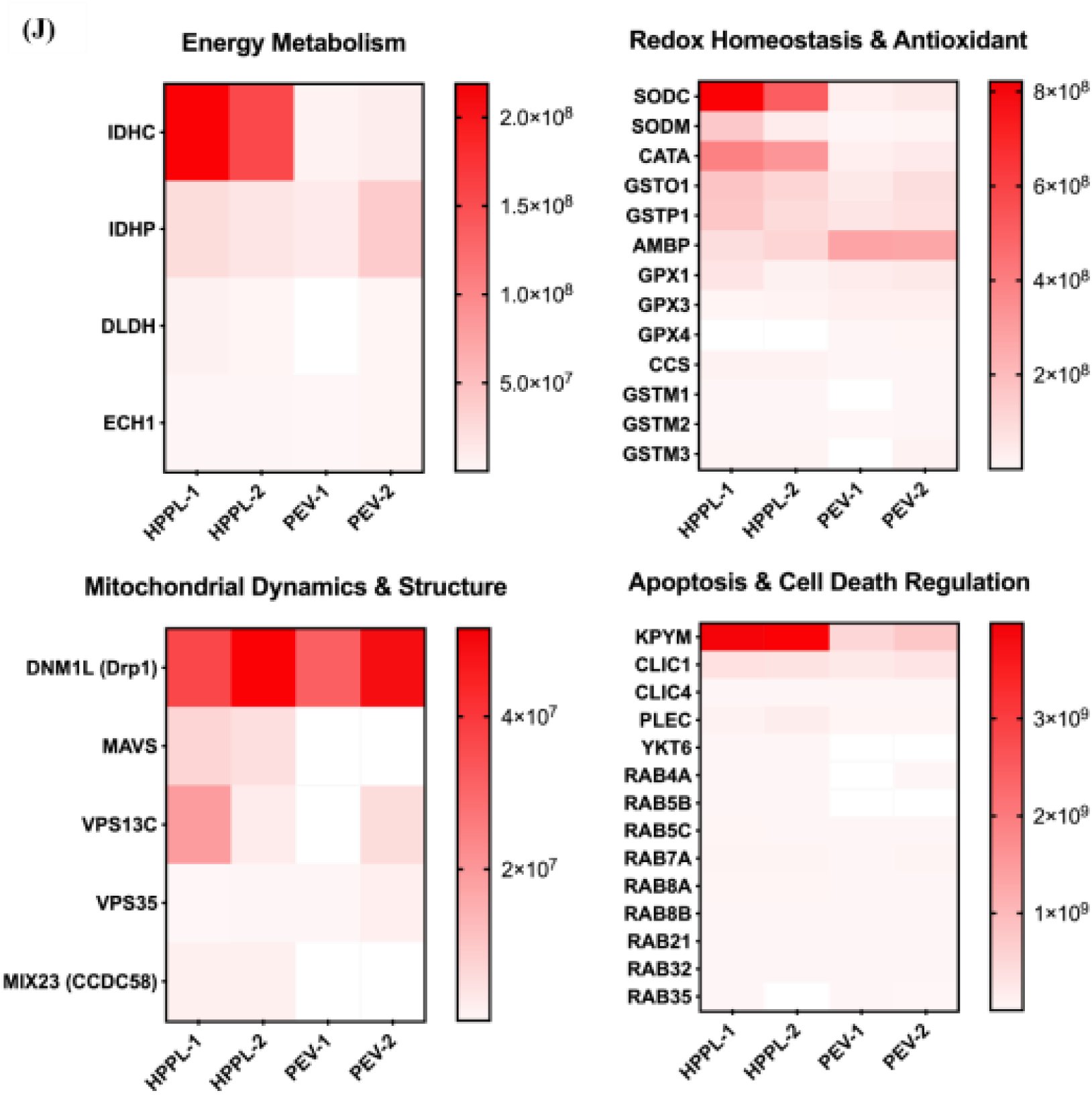
Biochemical and biophysical profiling of HPPL and PEVs. (A) Total protein concentration of HPPL and PEVs measured by BCA assay, n = 6 (B) Representative Cryo-EM image of PEVs, Scale bar: 100 nm. Black arrows represent the lipid bilayer membrane in PEVs (C) Average particle size of PEVs, as measured by DLS (D) Particle concentration (EV particles/mL) and size determination (nm) of PEVs analyzed by NTA n = 3 (E) Images displaying human-specific exosome antibody array after incubation with 3 pooled samples of PEVs showing EVs markers TSG101, ALIX, CD81, FLOT1, CD63, and ANXA5, cell adhesion protein ICAM and EpCAM, and the cis-Golgi marker GM130 (F) Quantification of EV markers relative to internal positive control (set as 100%). (G) SDS-Page of PEVs with Coomassie blue staining (H) Western blot analysis of PEVs for the presence of CD41, CD42a and CD62P, with β-actin used as an internal control. Uncropped western blot and exo-check membrane is shown in supplementary Figure 1 (I) Relative intensities of CD41, CD42a and CD62P were obtained by normalizing with the total β-actin level. Data are presented as mean ± SD, ****p < 0.0001 vs. PEVs; and ^###^p < 0.001, ^##^p < 0.01 ^#^p < 0.05 vs. MitoQ (J) Proteomic profiling heatmaps of mitochondrial-associated proteins in HPPL and PEVs represent the expression profiles of mitochondrial-related proteins, generated through a secondary analysis of proteomic datasets originally published by Delila L. *et al.* [32]. The data compare the protein compositions of HPPL and PEVs in regard with energy metabolism, redox homeostasis, mitochondrial dynamics, structure, apoptosis and cell death regulation.

### 3.2 Proteomic profiling reveals Mitochondrial-associated proteins in HPPL and PEVs

Our proteomic analysis of HPPL and PEVs identified a broad repertoire of mitochondrial-associated proteins involved in energy metabolism, apoptosis regulation, organelle dynamics, and redox homeostasis shown in Supplementary Table 1. Enzymes associated with mitochondrial energy metabolism included isocitrate dehydrogenase isoforms (IDHc/IDHp), enoyl CoA hydratase 1 (ECH1), and dihydrolipoamide dehydrogenase (DLDH), indicating the presence of tricarboxylic acid (TCA) cycle pathway, fatty acid β-oxidation, and aerobic respiration. Proteins implicated in apoptosis and mitochondrial membrane regulation were also detected, including BID, AIFM1, BAX, and cyclophilins (PPIA, PPIB, PPIF), suggesting enrichment of mitochondrial cell-death regulatory components. In addition, multiple cytoskeletal and membrane-trafficking proteins were identified, including CLIC1, CLIC4, and plectin (PLEC), which participate in mitochondrial-cytoskeletal interactions. Several Ras-related Rab proteins (RAB4A, RAB32, RAB5, RAB7, RAB8, RAB21, and RAB35) involved in mitochondrial dynamics and mitophagy, were also present, along with YKT6, a SNARE protein linked to autophagosome–lysosome fusion. Proteins linked to mitochondrial structural integrity and dynamics, including DNM1L (DRP1), MAVS, VPS13C, VPS35, and CCDC58, were also identified. Finally, proteins critical for redox regulation and antioxidant defense were abundantly detected, including superoxide dismutases (SOD1/2), catalase, glutathione peroxidases (GPX1/3/4), glutathione S-transferases, CCS, and AMBP, highlighting strong representation of oxidative stress modulating pathways in platelet-derived materials (Figure 2J).

### 3.2. Cellular Uptake of HPPL and PEVs in neuronal N2A Cells

To investigate the uptake and localization of HPPL and PEVs within neuronal cells, we assessed their intracellular uptake in N2A cells using Alexa Fluor-488-labeled HPPL and PEVs. Confocal microscopy revealed efficient cytoplasmic uptake of both fluorescently labeled PEVs and HPPL within 1 hr (Figure 3A & B). MitoTracker Red staining was used to visualize the mitochondrial network. Although limited colocalization was observed under baseline conditions using Pearson’s correlation coefficient of 0.33 for HPPL, the latter was found to be 0.68 for PEVs (Figure 3C & D), suggesting that PEVs have relatively higher partial spatial proximity with mitochondria.

**Figure 3.**
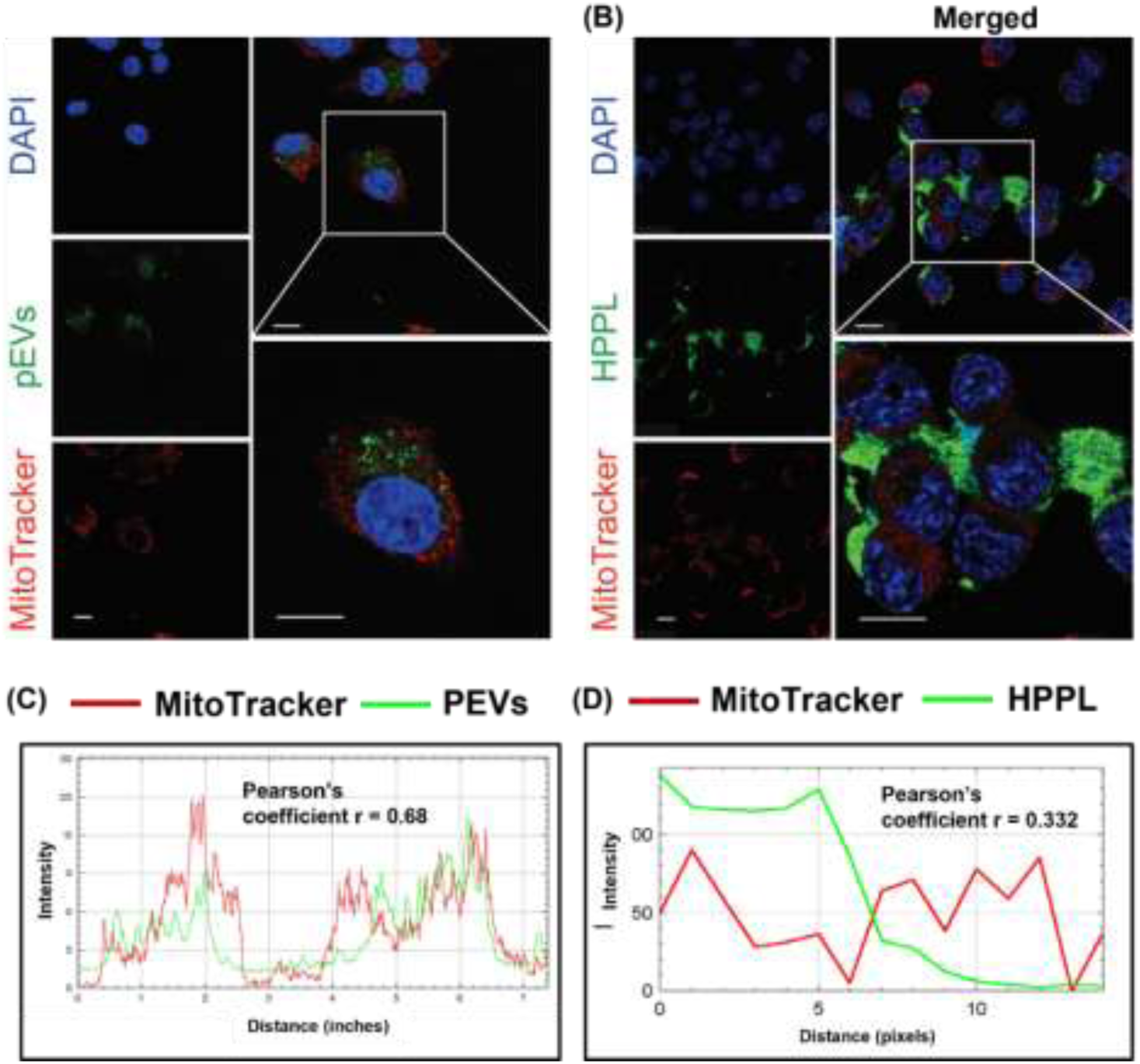
Cellular uptake of HPPL and PEVs in N2A neurons and their spatial relationship with mitochondria. (A) Cellular uptake of PEVs, Scale bar: 10 μm,100-fold magnification (B) Cellular uptake of HPPL, Scale bar: 10 μm, 40-fold magnification. (C) Quantified Pearson’s correlation coefficients were calculated to assess the degree of spatial overlap between green (PEVs) and red (MitoTracker) signals (D) Quantified Pearson’s correlation coefficients were calculated to assess the degree of spatial overlap between green (HPPL) and red (MitoTracker) signals using the Coloc2 plugin in ImageJ. Nuclei were stained with DAPI (blue), while HPPL or PEVs were labeled with AlexaFluor-488 (green), and MitoTracker (red) marked mitochondria.

### 3.3. HPPL and PEVs improve cell viability and confer neuroprotection against rotenone-induced complex I inhibition

The overall workflow of cellular experiments is presented in Figure 4A. To confirm the dose of HPPL, PEVs and rotenone to be used in the present study, pre-liminary dose-response experiments were conducted (Supplementary Figure 2). MitoQ, a mitochondria-targeted antioxidant derived from ubiquinone, was used as a positive antioxidant control at a dose that doesn’t impact cell viability (data not shown). Finally, 5 μM rotenone exposure for 24 hr was selected for the induction of mitochondrial dysfunction in both cells in all the subsequent experiments. N2A and SH-SY5Y cells were pre-treated with HPPL and PEVs for 1 hr prior to exposure to 5 µM rotenone. To assess cell survival, we employed the CCK-8 assay, which showed that exposure to rotenone reduced cell viability by an average of 61.98 % in N2A cells and 63.47 % in SH-SY5Y cells compared to their respective controls at 24 hr. MitoQ significantly restored cell viability by 89.77 % in N2A cells and 81.34 % in SH-SY5Y cells compared to the rotenone-treated group. Cells pre-treated with HPPL and PEVs for 1 hr before exposure to 5 μM rotenone demonstrated significant protection against rotenone-induced mitochondrial toxicity, specifically in N2A cells, HPPL increased cell survival by 58.04 % and PEVs by 43.62 % compared to the rotenone-only group (Figure 4B). Similarly, in SH-SY5Y cells, HPPL improved viability by 95.41 %, while PEVs enhanced survival by 59.96 % relative to rotenone treatment (Figure 4C). In parallel, cell viability was assessed in N2A cells using the Live/Dead assay, where viable cells showed green fluorescence and dead cells exhibited red fluorescence. Rotenone exposure significantly increased red fluorescence, indicating a higher proportion of dead cells. Pretreatment with HPPL or PEVs markedly attenuated rotenone-induced cell death, as evidenced by a visible reduction in red-stained cells, comparable to the protective effects observed with MitoQ (Figure 4D and E). These findings highlight the strong neuroprotective capacity of HPPL and PEVs against rotenone-induced cytotoxicity.

**Figure 4.**
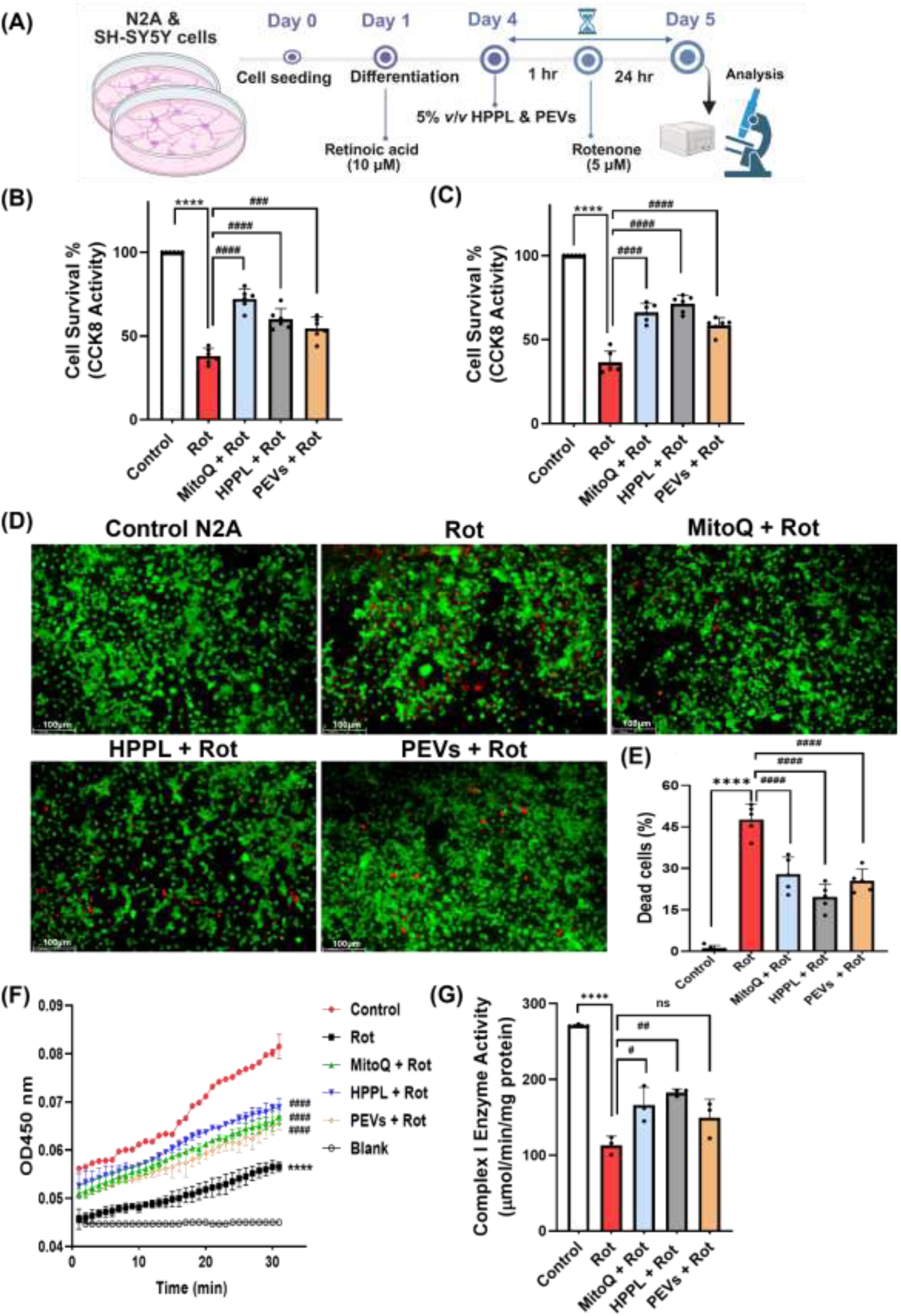
Neuroprotective effects of HPPL and PEVs on rotenone-induced mitochondrial dysfunction in neuronal cells. (A) Experimental design *in vitro*. (B) The effect of HPPL and PEVs in comparison to MitoQ treatment on the cell survival of N2A cells (n = 6) and (C) SH-SY5Y cells exposed to 5 μM rotenone (n = 6) (D) Representative fluorescence images from the live/dead cell assay showing green (live) and red (dead) N2A cells, scale bar: 100 μm (n = 6) (E) Quantification of dead cell percentage based on fluorescence image analysis from 5 random field areas (n = 5) (F) OD values per minute for mitochondrial complex I activity (G) Normalized complex I activity (μmol/min) with protein content (mg) in N2A cells (n = 3). Results expressed as a mean ± SD using one-way ANOVA comparisons tests; ****P < 0.0001 vs control; and ^###^P < 0.001, ^####^P < 0.0001 vs rotenone.

Next, to confirm whether these protective effects of HPPL and PEVs are linked to mitochondrial function, Complex I (NADH: ubiquinone oxidoreductase) enzymatic activity was evaluated in N2A cells following rotenone treatment. Rotenone exposure significantly inhibited Complex I activity by 58.23% within 24 hr compared to untreated control cells, confirming effective disruption of mitochondrial respiratory function. Pre-treatment with HPPL and PEVs restored Complex I enzyme activity by 61.36% and 32.16%, suggesting mitigation of rotenone-induced mitochondrial impairment (Figure 4F and G). These findings further confirm that HPPL and PEVs support mitochondrial health, potentially through mechanisms involving preservation or restoration of Complex I functionality.

### 3.4. HPPL and PEVs improve cellular bioenergetics and mitochondrial respiration in N2A Cells

In our investigation of cellular respiration in N2A cells exposed to rotenone, Figure 5A illustrates the significant drop of ATP level 24 hr following rotenone reaching 56.63 % as compared to the control group, indicating impaired mitochondrial activity. Pretreatment with HPPL restored ATP levels by 57.73 % and PEVs led to even higher ATP levels by 65.15 % relative to the rotenone-only group, indicating a protective effect on mitochondrial function. Subsequently, we assessed lactate content to evaluate glycolytic activity and metabolic alterations induced by rotenone and the potential protective effects of HPPL and PEVs. Rotenone exposure led to an increase in lactate levels by 109.65 % in N2A cells, indicative of a strong metabolic shift towards glycolysis compared to control cells. However, pretreatment with HPPL and PEVs exhibited a non-significant reduction by 35.09 % and 34.70 % in lactate levels compared to the rotenone-treated group, suggesting modest recovery of rotenone-induced metabolic stress (Figure 5 B). These findings suggest that both HPPL and PEVs confer mitochondrial protection and improve energy restoration under conditions of oxidative stress. Having observed these changes, we next investigated alterations in OXPHOS using the Seahorse MitoStress test. To avoid the confounding effect of direct complex I inhibition by rotenone, we instead used H₂O₂ as the inducer of oxidative stress. Treatment with H_2_O_2_ markedly reduced key OCR parameters, including basal respiration, ATP-linked respiration, maximal respiration, proton leak, and spare respiratory capacity, indicative of oxidative stress-induced mitochondrial dysfunction. These reductions confirmed that H_2_O_2_ effectively impairs mitochondrial bioenergetics, validating its use as a model to evaluate the protective effects of HPPL and PEVs (Figure 5C). Pre-treatment with HPPL, PEVs, and MitoQ significantly improved basal respiration compared with H_2_O_2_-treated cells (Figure 5D). Notably, HPPL and MitoQ also restored maximal respiration to a significant extent, whereas PEVs showed only a modest, non-significant effect (Figure 5E). In contrast, none of the treatments had a measurable impact on ATP-linked respiration and coupling parameters such as proton leak, or spare respiratory capacity (Figure 5F–H), suggesting that HPPL and PEVs primarily enhance mitochondrial bioenergetics in H_2_O_2_-treated cells by improving basal and maximal respiration rates.

**Figure 5.**
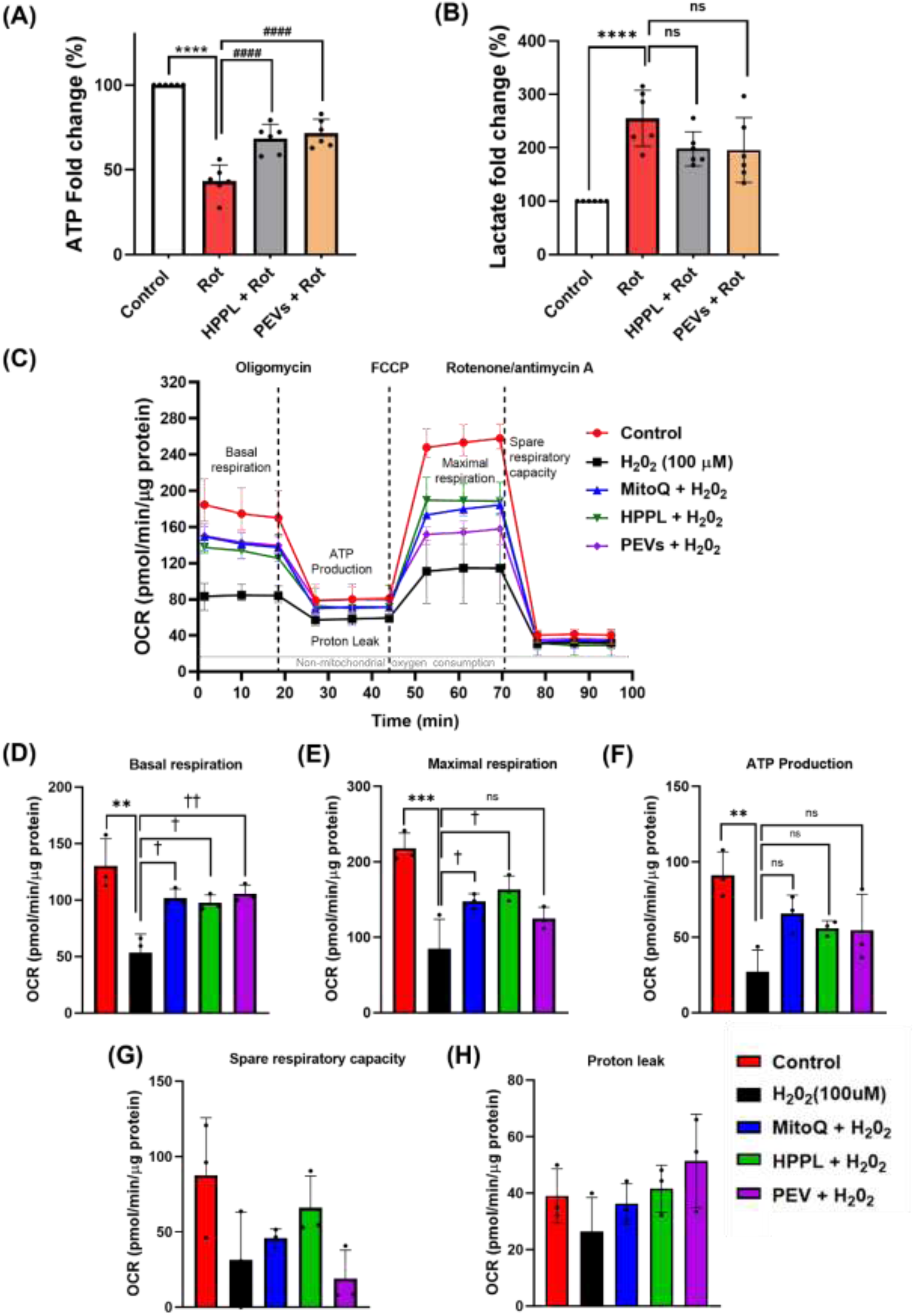
Effect of HPPL and PEVs on mitochondrial respiration and metabolic stress in N2A cells. (A) Measurement of intracellular ATP levels and (B) lactate levels as an indicator of glycolytic shift following treatment with HPPL or PEVs in N2A cells exposed to 5 μM rotenone. (C) Representative OCR trace from the Seahorse Mitostress assay in N2A cells treated with 100 μM for 24 hr, showing dynamic changes in respiration. (D-H) Bar graphs quantifying and comparing key OCR parameters, including (D) basal respiration, (E) maximal respiration, (F) ATP-linked respiration, (G) spare respiratory capacity and (H) proton leak following pre-treatment with HPPL, PEVs and MitoQ in H_2_O_2_-treated cells. Data represent the mean ± SD from three independent experiments using one-way ANOVA comparisons tests. *P < 0.05, **P < 0.01, ***P < 0.001, ****P < 0.0001 vs control; ^###^P < 0.001, ^####^P < 0.0001 vs rotenone; and ^†^P < 0.05, ^††^P < 0.01 vs H_2_O_2_.

### 3.5. Anti-oxidative potential of HPPL and PEVs against rotenone treated neuronal cells

To evaluate the antioxidative properties of HPPL and PEVs following rotenone exposure in cells, we first assessed intracellular ROS levels using the DCFDA fluorescent probe. Rotenone treatment markedly induces intracellular ROS generation as observed by elevated DCFDA fluorescence in both SH-SY5Y (Figure 6A) and N2A cells (Figure 6B). Pretreatment with HPPL and PEVs significantly reduced DCFDA signal intensity, suggesting an attenuation of general oxidative stress. Quantification of DCFDA mean fluorescence intensity is shown in Figure 6E and F. To further confirm mitochondrial-specific oxidative stress, we employed MitoSOX Red staining, which selectively detects mitochondrial superoxide generation. Rotenone exposure led to a strong increase in MitoSOX red fluorescence, consistent with elevated mitochondrial ROS generation. This effect was markedly reduced in cells pretreated with HPPL, supporting their protective role in limiting mitochondrial oxidative damage (Figure 6C). However, PEVs pretreatment did not significantly reduce MitoSOX fluorescence, suggesting a less pronounced effect on mitochondrial superoxide levels compared to HPPL (Figure 6D). Quantification of MitoSOX fluorescence intensity is shown in Figure 6G and H. In addition, we measured GPx activity as a marker of antioxidant defense. GPx activity decreased by 57.37% in the rotenone-only group compared to control cells, indicating impaired enzymatic defense against oxidative stress. However, pretreatment with HPPL and PEVs significantly restored GPx activity by 60.71% and 53.08%, respectively, relative to rotenone-treated cells (Figure 6I). Additionally, to confirm the overall antioxidative potential of HPPL and PEVs preparations, we employed the ORAC assay, normalized to protein content. Both HPPL and PEVs demonstrated higher antioxidant activity, exhibiting 126.93% and 216.84% activity per mg protein relative to the positive control MitoQ. In contrast, ATP and plasma (negative controls) showed lower antioxidant capacity (Figure 6J). Together, these findings establish that HPPL and PEVs effectively reduce both intracellular and mitochondrial ROS, restoring enzymatic antioxidant defense, and ultimately confirming their strong radical-scavenging potential through ORAC analysis.

**Figure 6.**
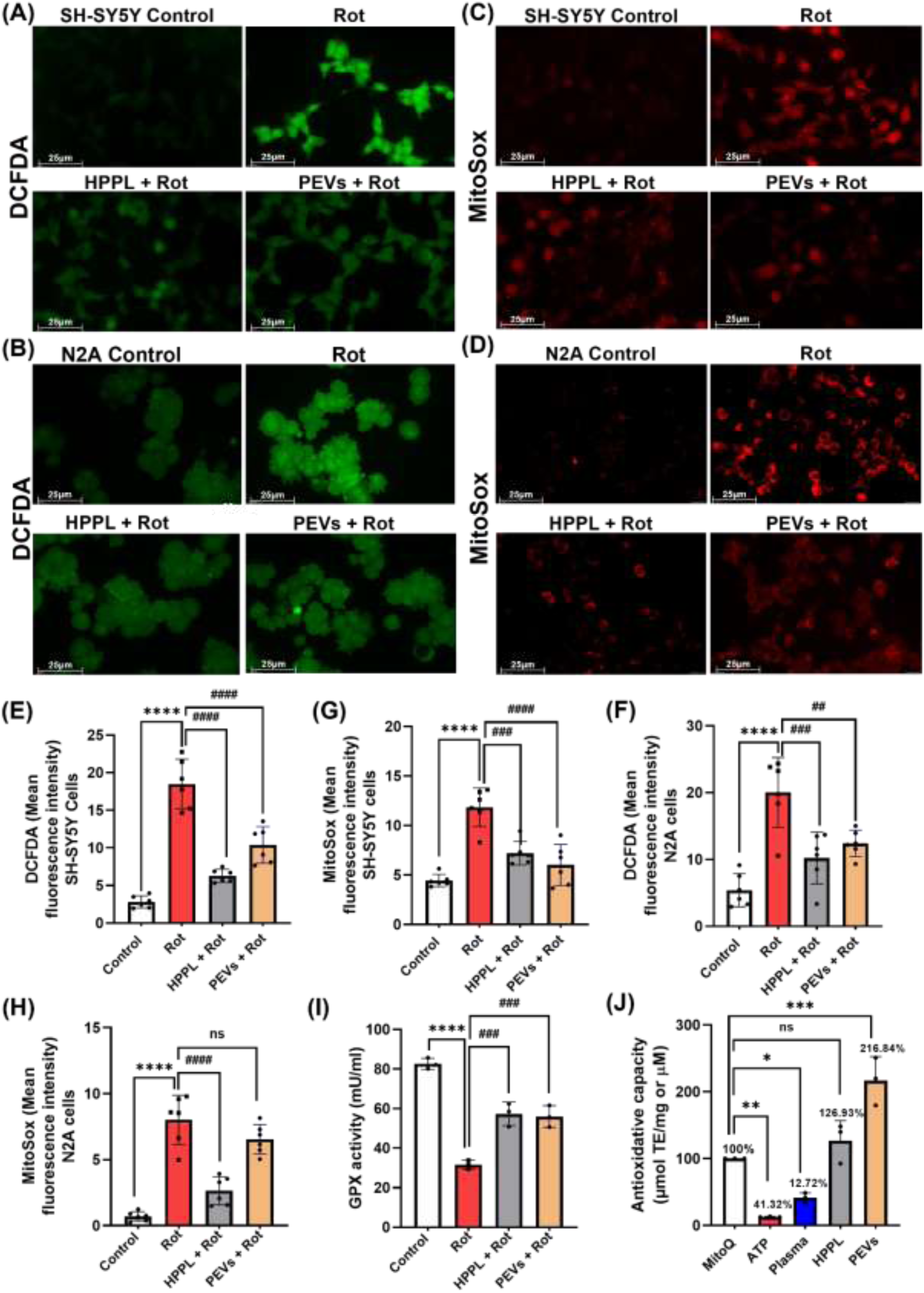
Effect of HPPL and PEVs on the oxidative stress. (A) Intracellular ROS depicted by DCFDA staining in SH-SY5Y cells, scale bar 25 μm, n = 6 (B) Intracellular ROS in N2A cells, scale bar 25 μm, n = 6 (C) Mitochondrial ROS depicted by mitoSox staining in SH-SY5Y cells, scale bar 25 μm, n = 6 (D) MitoSox staining in N2A cells, scale bar 25 μm, n = 6. All representative images were captured using advance fluorescence microscopy at 40-fold magnification (E) Mean fluorescence intensity quantification of DCFDA staining in SH-SY5Y cells (F) Quantification of DCFDA staining in N2A cells (G) Mean fluorescence intensity quantification of mitoSox staining in SH-SY5Y cells (H) Quantification of mitoSox staining in N2A cells. Mean intensities were measured from 6 different fields for each group (I) GPX activity in N2A cells (J) Antioxidant capacity of HPPL and PEVs in triplicate is expressed in µmol Trolox equivalents (TE) per mg total protein content and MitoQ was used as a positive control (100%). Values are expressed as mean ± SD (n = 3) one-way ANOVA comparisons tests; ****P < 0.0001, ***P < 0.001, **P < 0.01, *P < 0.05 vs control, MitoQ; and ^####^P < 0.0001, ^###^P < 0.001, ^##^P < 0.01, ^#^P< 0.05 vs rotenone group.

### 3.6. HPPL and PEVs maintain mitochondrial integrity and membrane potential

JC-1 staining was used to evaluate mitochondrial ΔΨm, a critical indicator of mitochondrial function. Under physiological conditions, healthy mitochondria exhibit high ΔΨm, allowing JC-1 to form red-emitting aggregates. Rotenone-treated live cells showed a marked increase in green fluorescence, indicating mitochondrial depolarization and loss of ΔΨm. In contrast, cells pretreated with HPPL and PEVs maintained a higher red-to-green fluorescence ratio, demonstrating preserved ΔΨm and mitochondrial function (Figure 7A). To further assess mitochondrial function, MitoTracker Red staining was performed. In untreated control cells, MitoTracker Red strongly accumulated in active mitochondria, indicating preserved mitochondrial mass and membrane integrity. In contrast, rotenone-exposed cells exhibited a marked decrease in MitoTracker Red mean fluorescence intensity, reflecting compromised mitochondrial integrity (Figure 7B). Notably, quantitative analysis of JC-1 (Figure 7C & D) and MitoTracker Red (Figure 7E) showed that pre-treatment with HPPL and PEVs effectively preserved ΔΨm and mitochondrial function under oxidative stress conditions. Next, TEM provided high-resolution images of mitochondrial ultrastructure, revealing morphological changes associated with rotenone exposure, including mitochondrial swelling, disrupted cristae, and outer membrane rupture, hallmarks of mitochondrial damage. Conversely, cells pretreated with HPPL and PEVs displayed mitochondria structure with intact membranes and well-defined cristae, indicating preserved mitochondrial structure (Figure 7F-H). Together, these findings confirm that HPPL and PEVs help preserve mitochondrial integrity, structure, and function by mitigating the deleterious effects of rotenone-induced mitochondrial stress.

**Figure 7.**
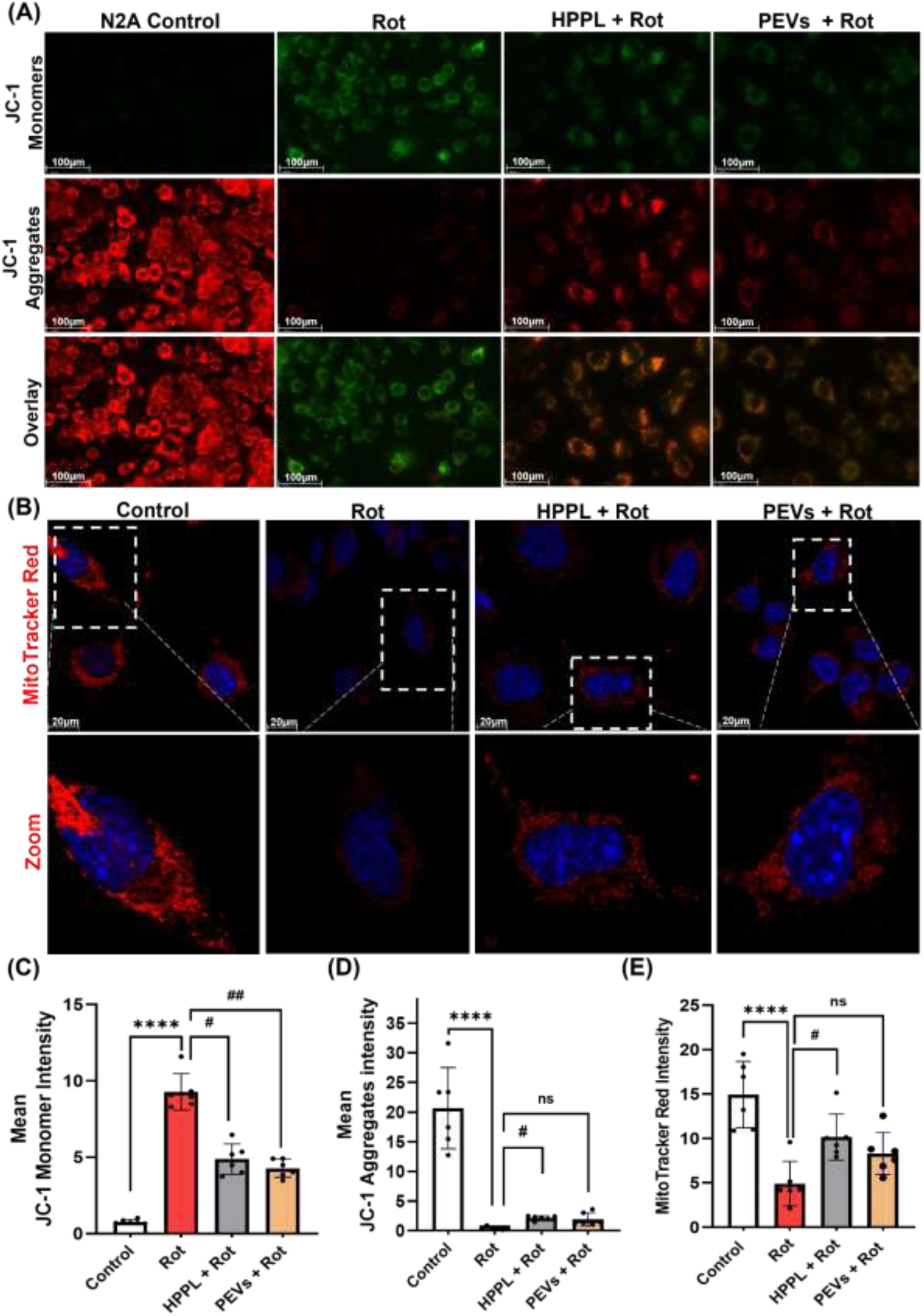
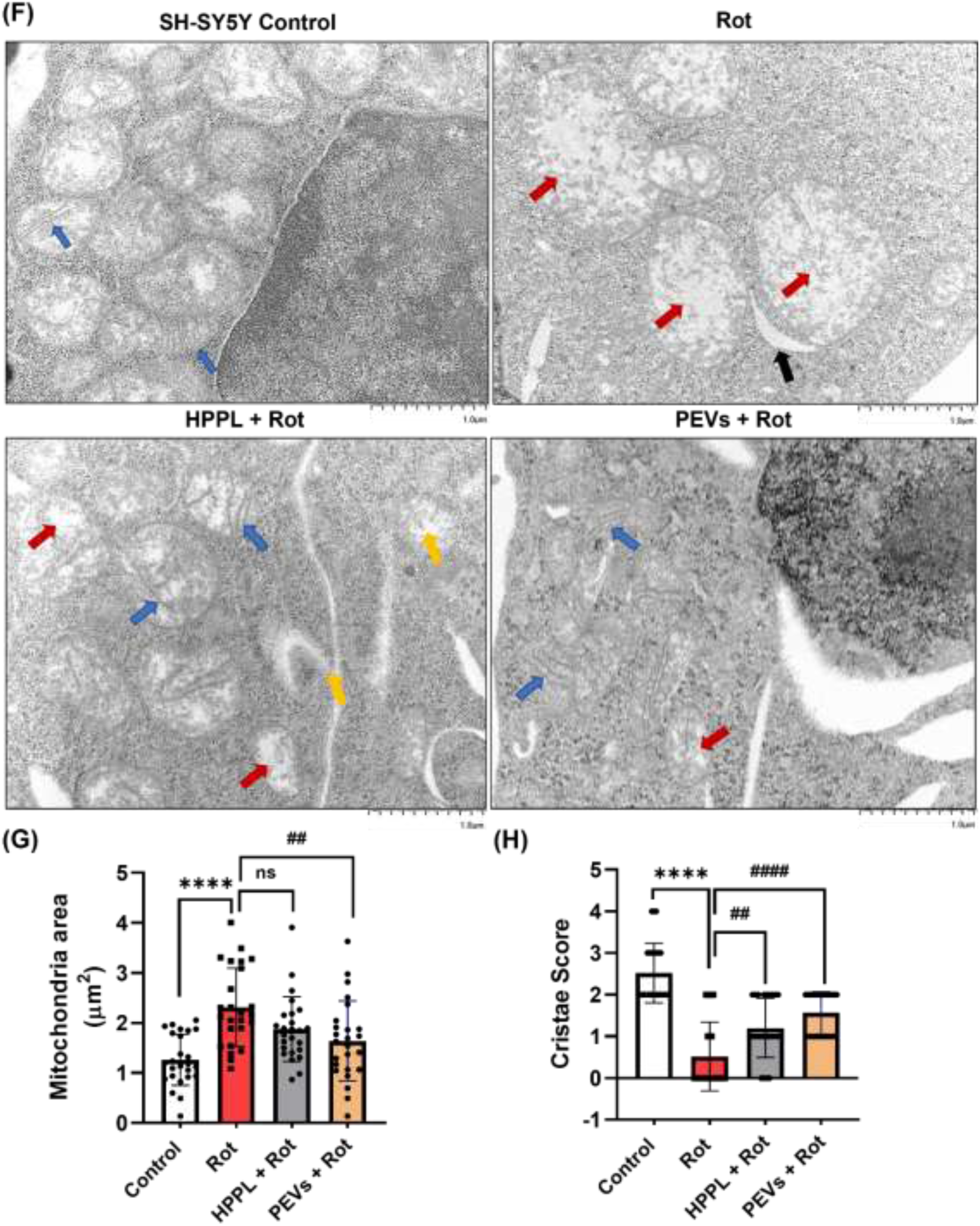
Effect of HPPL and PEVs on mitochondrial structure and function. (A) mitochondrial membrane potential assessed by JC-1 fluorescent probe captured using advance fluorescence microscopy at 40-fold magnification; scale bar: 100 μM (B) mitochondrial permeability assessment by MitoTracker red staining and counterstained with DAPI to visualize mitochondria viewed under confocal microscope, 20 μM, 63-fold magnification (C) Quantification of JC-1 monomer mean fluorescence intensity (D) Mean fluorescence intensity quantification of JC-1 aggregation (E) Mean fluorescence intensity quantification of MitoTracker red measured from 6 different fields from 3 independent experiments for each group (F) Ultrastructure of SH-SY5Y cells mitochondria observed under TEM, representative images captured at 12-fold magnification, scale bar: 1 μm. Blue arrows represent organized dense mitochondrial cristae, red arrows show collapsed mitochondrial matrix with abnormal cristae, black arrow defines mitochondrial membrane integrity and yellow arrow shows damaged mitochondrial membranes. (G) Quantification of mitochondrial area (um^2^) in SH-SY5Y cells (H) Measurement of the cristae quality score from 25 mitochondria from two biological replicates. Values are expressed as mean ± SD using one-way ANOVA comparisons tests; ****P < 0.0001, ***P < 0.001 vs control; ^####^P< 0.0001, ^###^P< 0.001, ^##^P< 0.01, ^#^P< 0.05 vs rotenone group.

### 3.7. HPPL and PEVs Promote Mitochondrial Biogenesis and fusion in neurons

Mitochondrial biogenesis is a vital adaptive process that sustains mitochondrial function and cellular energy homeostasis, particularly under conditions of oxidative stress. To assess the impact of rotenone-induced toxicity and the potential therapeutic effects of HPPL and PEVs, we evaluated the expression of PGC-1α, a key transcriptional coactivator and master regulator of mitochondrial biogenesis. Immunofluorescence analysis revealed that PGC-1α fluorescence was predominantly localized to the nucleus in control cells, showing high intensity consistent with active transcriptional function. Rotenone exposure led to a notable decrease in overall PGC-1α fluorescence intensity, particularly within the nuclear compartment. Interestingly, pretreatment with PEVs led to a pronounced increase in PGC-1α signal intensity predominantly in the nucleus, whereas HPPL treatment enhanced fluorescence in both the nucleus and cytoplasm. This differential distribution suggests that PEVs may preferentially activate nuclear transcriptional regulation, while HPPL promotes a broader upregulation and redistribution of PGC-1α, consistent with a more extensive enhancement of the mitochondrial biogenic response (Figure 8A and B). ELISA-based quantification further supported these findings as PGC-1α protein levels were significantly reduced by 41.28% in rotenone-treated cell lysates compared to untreated controls, indicating suppression of mitochondrial biogenic signaling. In contrast, cells pretreated with HPPL and PEVs exhibited a marked restoration of PGC-1α levels by 20.08% and 52.06% compared to rotenone-treated cells, suggesting increased upregulation of mitochondrial biogenesis by PEVs through activation of upstream regulatory mechanisms (Figure 8C). To further investigate mitochondrial dynamics, we examined the balance between DRP1, a key regulator of mitochondrial fission, and MFN1, an essential mediator of mitochondrial fusion. In control cells, MFN1 expression was prominent while DRP1 levels remained basal, indicating a balanced fusion and fission state with intact mitochondrial network organization. In contrast, rotenone treatment markedly increased DRP1 expression and significantly reduced MFN1 levels, reflecting a shift toward excessive mitochondrial fission and network fragmentation. Following HPPL and PEVs treatment, moderate reduction in DRP1 expression accompanied by pronounced recovery of MFN1 levels was observed, indicating re-establishment of mitochondrial dynamic equilibrium (Figure 8D and F). Next to determine the mitochondrial structural integrity, MFN1 co-localization with the outer mitochondrial membrane marker TOMM20 was further examined. In control cells, MFN1 showed widespread distribution and strong spatial overlap with TOMM20, consistent with intact mitochondrial fusion activity and network integrity. Rotenone exposure markedly disrupted this co-localization, indicating impaired fusion and increased mitochondrial fragmentation. Remarkably, in PEVs treated cells, MFN1 expression was visibly restored and showed enhanced spatial overlap with TOMM20 and HPPL expressed a moderate increase in MFN-1 levels, suggesting re-establishment of mitochondrial fusion machinery and preservation of mitochondrial architecture. TOMM20 intensity was also improved in these groups, further reflecting rescue of mitochondrial mass and membrane structure (Figure 8E and G). Together, the combined upregulation of PGC-1α, the coordinated normalization of DRP1 and MFN1 expression, and concomitant restoration of MFN1–TOMM20 co-localization highlights a comprehensive recovery of mitochondrial biogenesis and mitochondrial fusion–fission balance, a key aspect of mitochondrial quality control.

**Figure 8.**
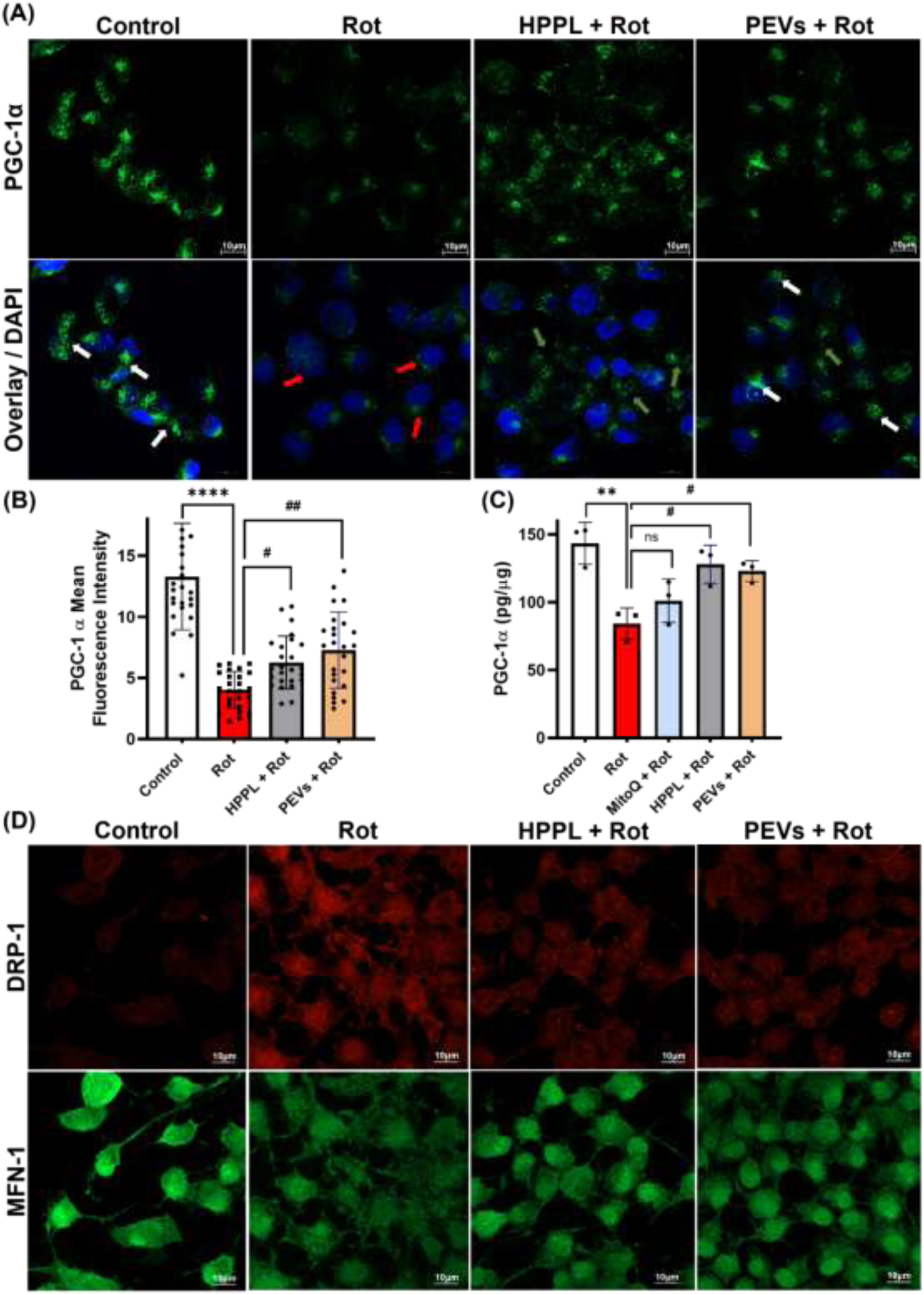
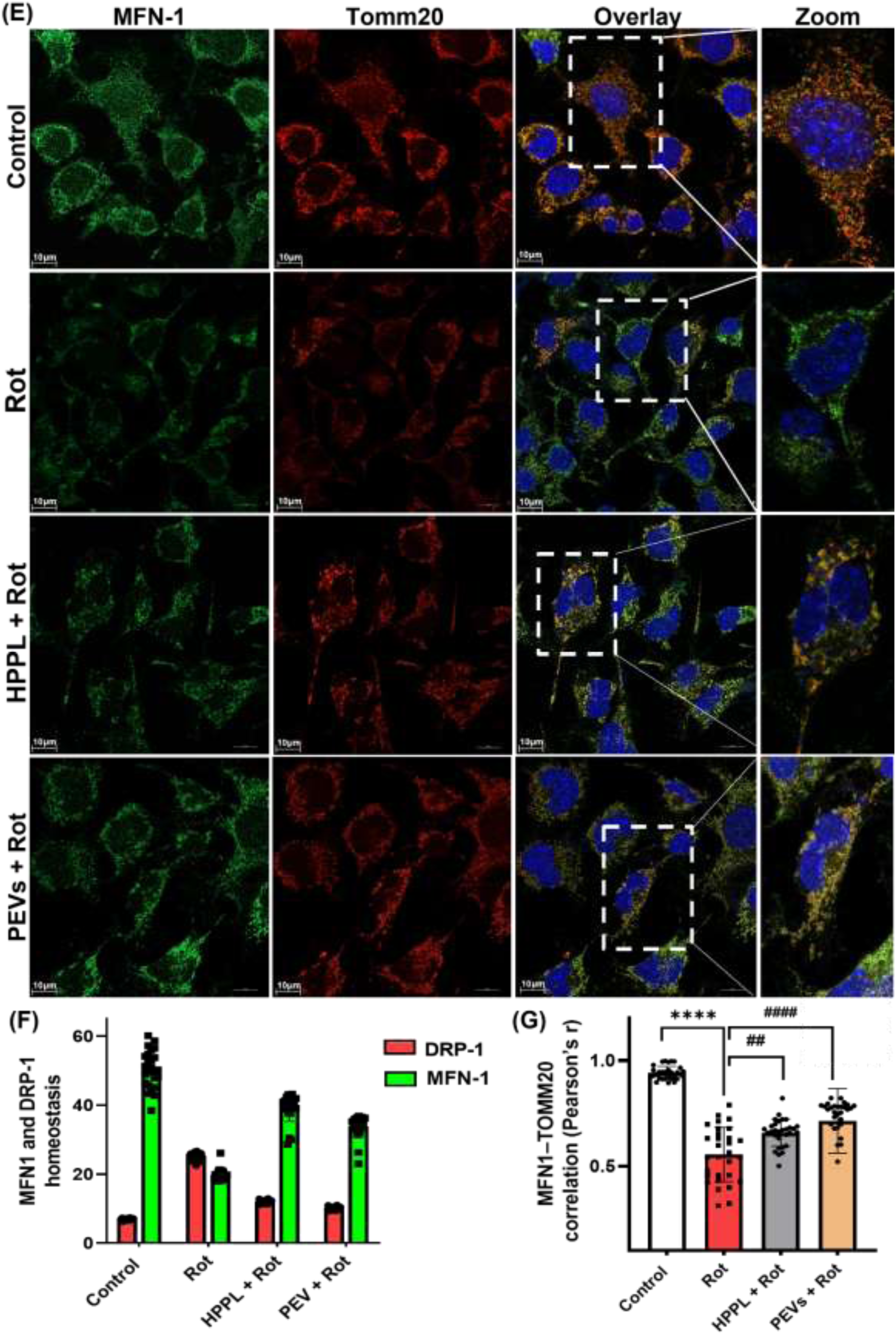
Impact of HPPL and PEVs on mitochondrial markers to assess mitochondrial biogenesis and fusion. (A) Representative immunofluorescence images showing PGC-1α, white arrows depict strong functional nuclear fluorescence, red arrows shows reduced overall fluorescence intensity, green arrows shows distinct fluorescence intensity in cytoplasm, Scale bar 10 μm, 63-fold magnification (B) Quantification of immunofluorescence signal intensity from 25 individual mitochondria from random fields across treatment groups (C) PGC-1α protein levels determined by ELISA in N2A cells across different treatment groups (D) Representative immunofluorescence images showing MFN1 and DRP1 expression in N2A cells across treatment groups. Scale bar = 10 µm; 63-fold magnification. (E) Representative images showing co-localization of MFN1 and TOMM20 in N2A cells, Scale bar 10 μm, 63-fold magnification (F) Quantification of MFN1 and DRP1 fluorescence intensity across treatment groups, analyzed from 25 individual mitochondria per group selected from random fields (G) Colocalization between MFN1 and TOMM20 shown using Pearson’s correlation coefficient (r), calculated by Coloc-2 plugin in ImageJ (Fiji). Values are expressed as mean ± SD (n = 25); ****P < 0.0001, **P < 0.01 vs control; and ^####^P < 0.0001, ^##^P < 0.01, ^#^P< 0.05 vs rotenone.

### 3.8. Mitochondrial Proteome Remodeling by HPPL and PEVs in N2A cells under Rotenone Stress

To elucidate the impact of HPPL and PEVs on rotenone-induced mitochondrial dysfunction, we performed comparative mitochondrial proteomic profiling in N2A cells across all experimental groups. This analysis revealed both shared and treatment-specific protein expression patterns. Using Ingenuity Pathway Analysis (IPA) (Supplementary figure 3), we examined canonical pathways related to mitochondrial function in N2A cells treated with rotenone, with or without pretreatment using HPPL or PEVs (Figure 9A). Rotenone treatment alone induced significant mitochondrial dysfunction, characterized by the inhibition of complex I consistent with rotenone’s known mechanism as a complex I inhibitor and the activation of complex III, cytochrome C and complex IV. The release of cytochrome c into the cytosol, triggered by these disruptions, ultimately promoted apoptosis in the rotenone-treated group.

**Figure 9.**
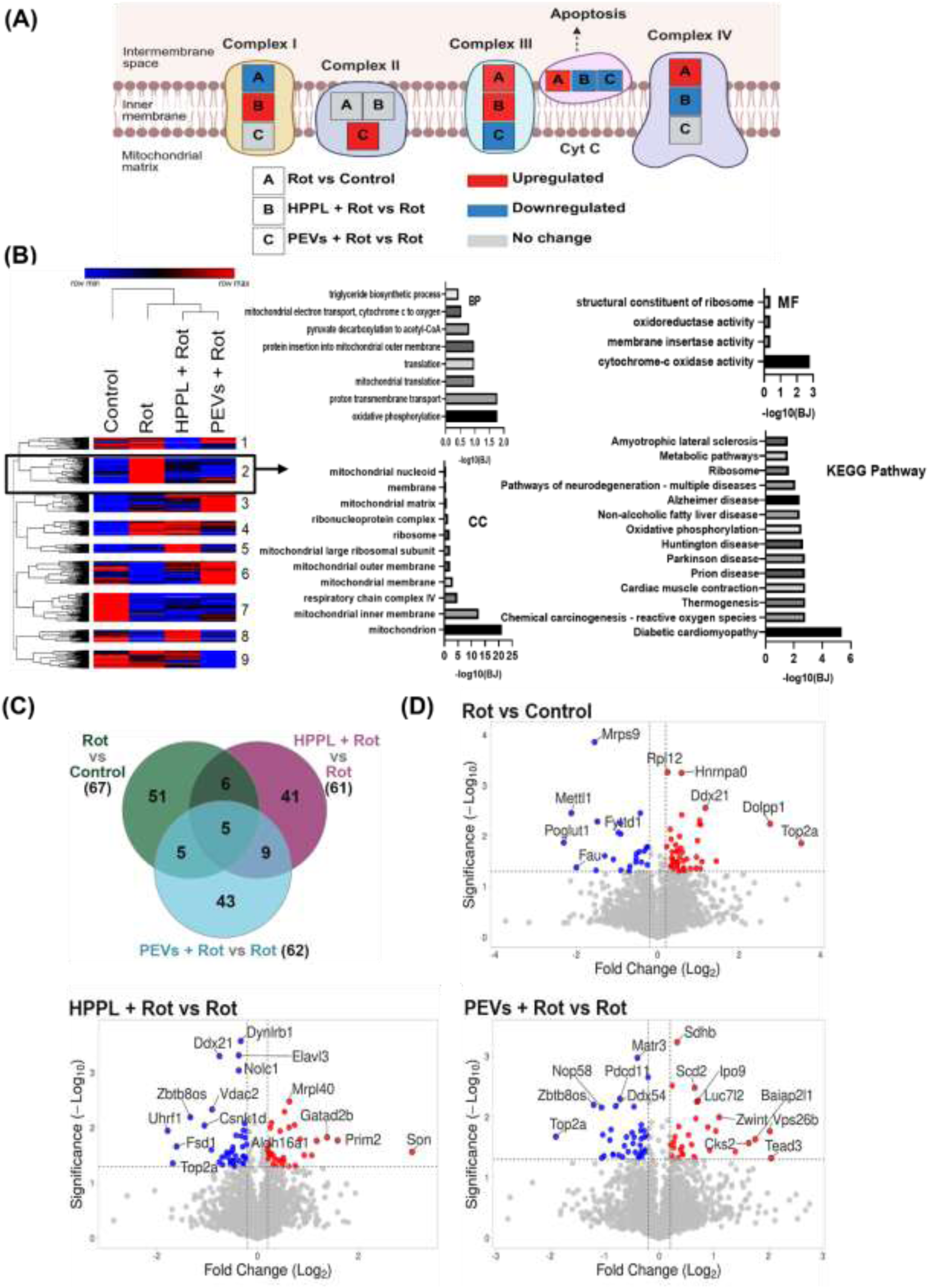
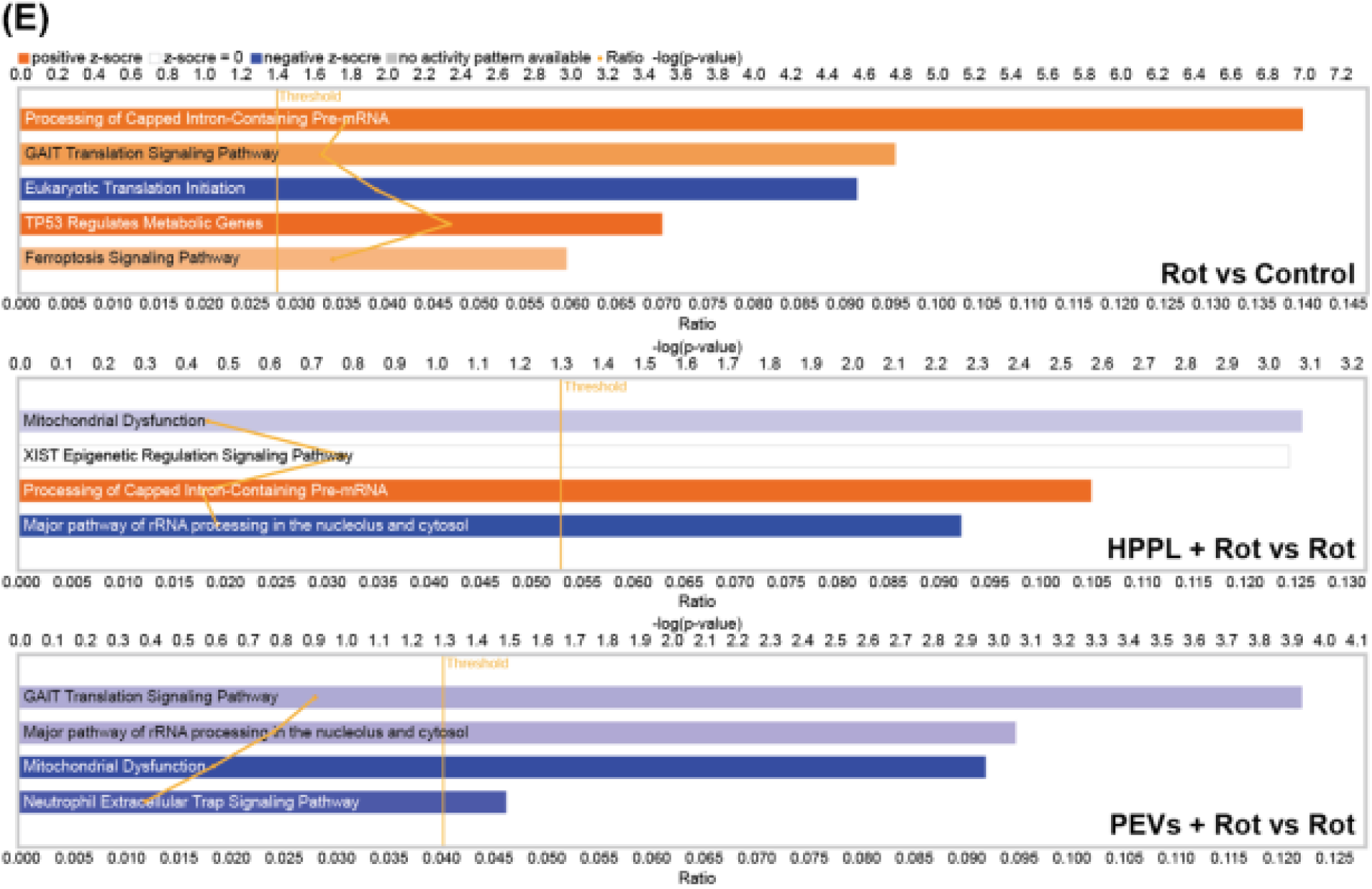
Mitochondrial proteomic profiling of N2a cells comparison of rotenone against control, HPPL + rotenone, and PEVs + rotenone against rotenone treatments. (A) Schematic reconstruction of ETC complex alterations inferred from IPA deep canonical pathways. Red: upregulation; blue: downregulation; grey: no significant change. (B) Heatmap of DEPs with log2 fold change (Log2FC) values, alongside GO (BP, MF, CC) and KEGG pathway enrichment. (C) Venn diagram showing shared and treatment-specific mitochondrial proteins among control, HPPL, and PEVs + rotenone groups. (D) Volcano plot of differentially expressed mitochondrial proteins (DEPs) comparing rotenone with control and HPPL or PEVs treated cells to rotenone group (p ≤ 0.05, fold change ≥ 1.2) (E) IPA canonical pathway analysis showing altered mitochondrial signaling networks (p < 0.05). Pathways color-coded by activation z-score: orange = activation, blue = inhibition, white = no change; green line indicates the ratio of DEPs per pathway.

Pretreatment with HPPL before rotenone exposure partially protected complex I–associated protein expression that was reduced by rotenone and was accompanied by modulation of downstream respiratory chain components. Notably, HPPL pretreatment attenuated cytochrome c–associated apoptotic signaling and reduced the activation of pathways linked to mitochondrial stress responses. In contrast, PEVs pretreatment produced a distinct mitochondrial response characterized by increased representation of complex II-related proteins together with reduced activation of complex III and cytochrome c-associated pathways. These findings suggest that HPPL and PEVs pre-treatment mitigates rotenone-induced mitochondrial dysfunction through partially distinct mechanisms: HPPL primarily restores complex I-related bioenergetic pathways, whereas PEVs appear to promote compensatory respiratory adaptations involving complex II while suppressing apoptotic signaling.

Heatmap analysis of the total proteome based on z-score normalized expression values identified nine modules (Figure 9B). Among these, module 2 (supplementary table 2) was selected for further analysis due to its biological relevance. In this module, protein expression levels were markedly increased in the rotenone-treated group but returned toward control levels following pretreatment with either HPPL or PEVs. Gene Ontology (GO) enrichment analysis revealed that these proteins were associated with oxidoreductase activity and cytochrome-c oxidase activity in the molecular function (MF) category, while cellular component analysis indicated localization within the mitochondrial matrix, membrane, nucleoid, and mitochondrion. Biological process enrichment further linked these proteins to mitochondrial electron transport (cytochrome c to oxygen), mitochondrial translation, and OXPHOS, consistent with KEGG pathway analysis.

To further characterize treatment-specific mitochondrial proteome alterations, we examined overlapping and DEPs using Venn diagrams (Figure 9C) and volcano plots (Figure 9D). Comparison of rotenone-treated cells with controls identified 67 mitochondrial proteins specifically altered by rotenone. Of these, 11 proteins overlapped with those observed in the HPPL + Rot vs. Rot comparison, while 50 proteins were unique to the HPPL pretreatment group, indicating substantial remodeling of the mitochondrial proteome following HPPL exposure. Similarly, 10 proteins overlapped between the Rot vs. control and PEVs + Rot vs. Rot comparisons, whereas 52 proteins were unique to the PEVs pretreatment group, suggesting that PEVs also induce distinct mitochondrial proteomic adaptations. Notably, 14 proteins were commonly regulated in both the HPPL + Rot vs. Rot and PEVs + Rot vs. Rot comparisons, indicating a subset of shared mitochondrial responses to platelet-derived treatments (Supplementary table 3). Volcano plot analysis further illustrated the differentially expressed mitochondrial proteins across the comparisons, highlighting both overlapping and treatment-specific proteomic responses involved in the mitigation of rotenone-induced mitochondrial stress (Supplementary table 4). IPA network analysis (Figure 9E) further highlighted the involvement of these proteins in mitochondrial stress signaling.

### 3.9. PEVs Mitigate Rotenone-Induced Developmental Toxicity in ZF Embryos

Schematic representation of the overall workflow for the ZF embryo experiments is shown in Figure 10A. To establish ZF as a model for assessing the *in vivo* biological effects of HPPL and PEVs, we optimized treatment concentrations using the zebrafish embryo toxicity (ZFET) assay (Supplementary Figure 4). For the safe and effective dosing conditions in ZF embryos, both HPPL and PEVs concentrations were optimized based on protein content and particle load. Preliminary assays (Supplementary Figure 5) identified that protein concentrations exceeding 30 µg/mL induced developmental toxicity. Therefore, HPPL and PEVs stocks were diluted to yield final exposure concentrations of approximately 20-25 µg/mL, respectively, both within the tolerated range. Additionally, high particle counts in undiluted PEVs were adjusted to avoid excessive exposure (∼10^9^ – 10^10^ particles/mL), ensuring biocompatibility. Exposure to 200 nM rotenone resulted in a marked decline in embryo survival (Figure 10B) and a significant delay in hatching (Figure 10C) from 4 hpf to 96 hpf, indicating developmental toxicity with multiple morphological defects, including head and tail malformations, delayed body axis elongation, pericardial edema, and impaired motility. In contrast, pretreatment with PEVs showed pronounced improvement in both survival rates and hatching time and significantly fewer abnormalities, restoring them to levels comparable with untreated controls however, HPPL showed only partial improvement, suggesting PEVs are more effective than HPPL in mitigating rotenone-induced developmental toxicity (Figure 10D and E, supplementary figure 6). These findings highlight the superior *in vivo* efficacy of PEVs in promoting embryonic resilience against oxidative mitochondrial stress.

**Figure 10.**
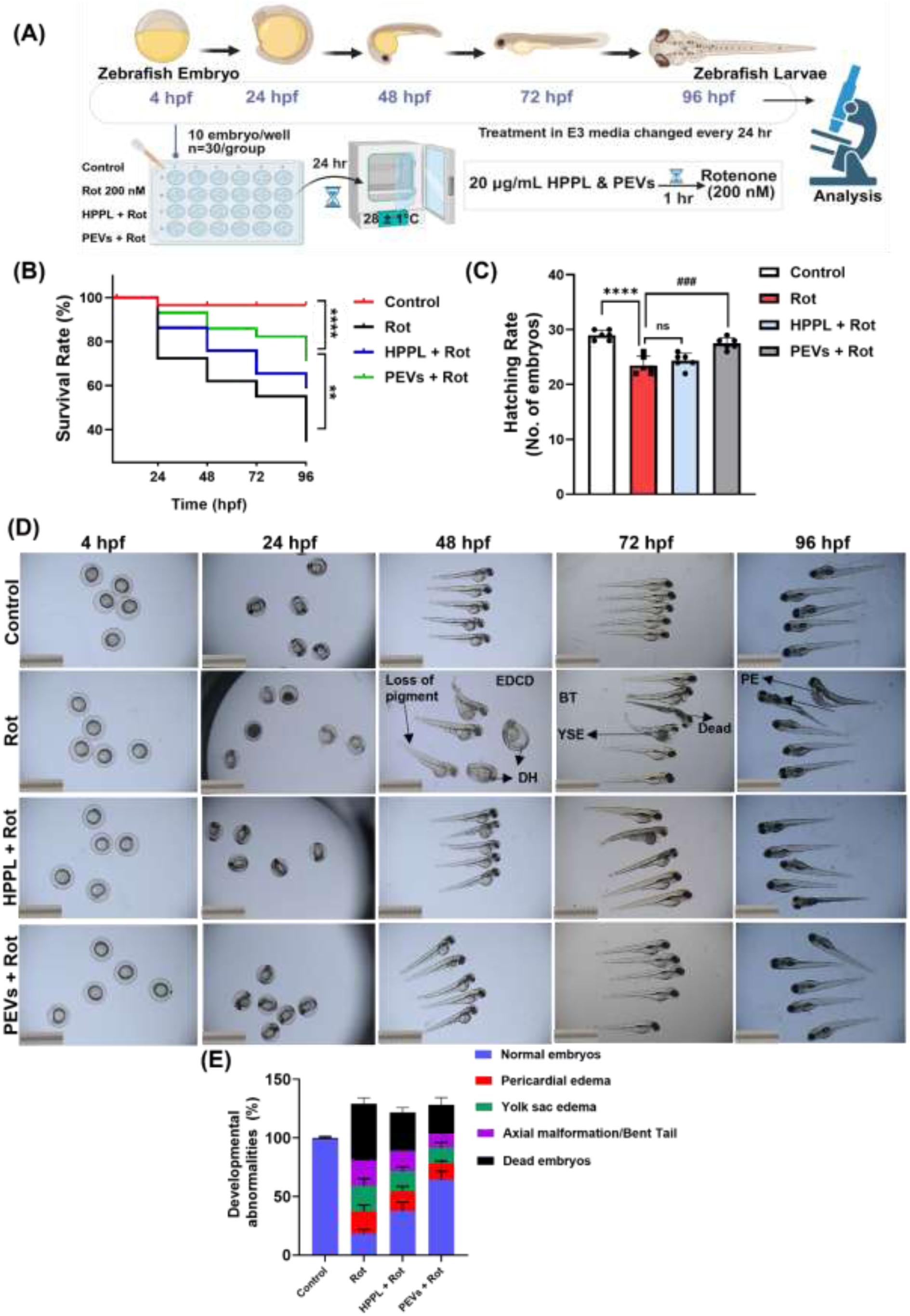
HPPL and PEVs improve survival and development in rotenone-exposed ZF embryos. (A) Experimental design illustrating the timeline of ZF embryo exposure to rotenone and co-treatment with HPPL and PEVs (n = 30/group/experiment) (B) Effect of HPPL and PEVs on survival rate of ZF embryos exposed to rotenone assessed every 24 hr (C) Effect of HPPL and PEVs on hatching rate at 48 hpf from six biological replicates (D) Representative morphological images of ZF embryos from each group at 4, 24, 48, 72, 96 hpf (E) Quantification of developmental abnormalities including normal embryos, PE, YSE, AM/BT, DE (% affected embryos per group), compiled from six independent experiments. EDCD: Embryos died with circulatory defects; PE: Pericardial edema; DH: Delayed hatching; YSE: Yolk sac edema; AM: Axial Malformation, BT: Bent tail. Values are expressed as mean ± SD (n = 6); ****P < 0.0001 vs control; ^###^P < 0.001 vs rotenone treated embryos

### 3.10. Mitochondrial targeting in ZF embryo and biodistribution of HPPL and PEVs

To assess biodistribution and mitochondrial uptake in whole embryos, we labeled both HPPL and PEVs with Alexafluor-488 and tracked their colocalization with mitochondria co-stained using MitoTracker red. Fluorescent imaging revealed the distribution of HPPL and PEVs mainly in key developmental regions of ZF embryos, indicating efficient uptake and systemic distribution (Figure 11A). Notably, at 24 hpf, the labeled HPPL and PEVs were largely unable to penetrate the protective chorion of the embryo, however, subsequent imaging showed that they successfully entered the embryo after this time point, suggesting time-dependent internalization. This indicates that HPPL and PEVs can reach various tissues within the embryo and interact with multiple cellular systems during embryonic development. Moreover, a visible overlap between the PEVs signal and MitoTracker red indicates successful mitochondrial uptake. Colocalization analysis revealed a high Pearson’s coefficient of 0.93 for PEVs, indicating that a substantial portion of the fluorescent signal was localized within mitochondria, suggesting effective mitochondrial targeting. In contrast, HPPL showed minimal spatial association with mitochondria, with a Pearson’s coefficient of 0.34 (Figure 11 B & C).

**Figure 11.**
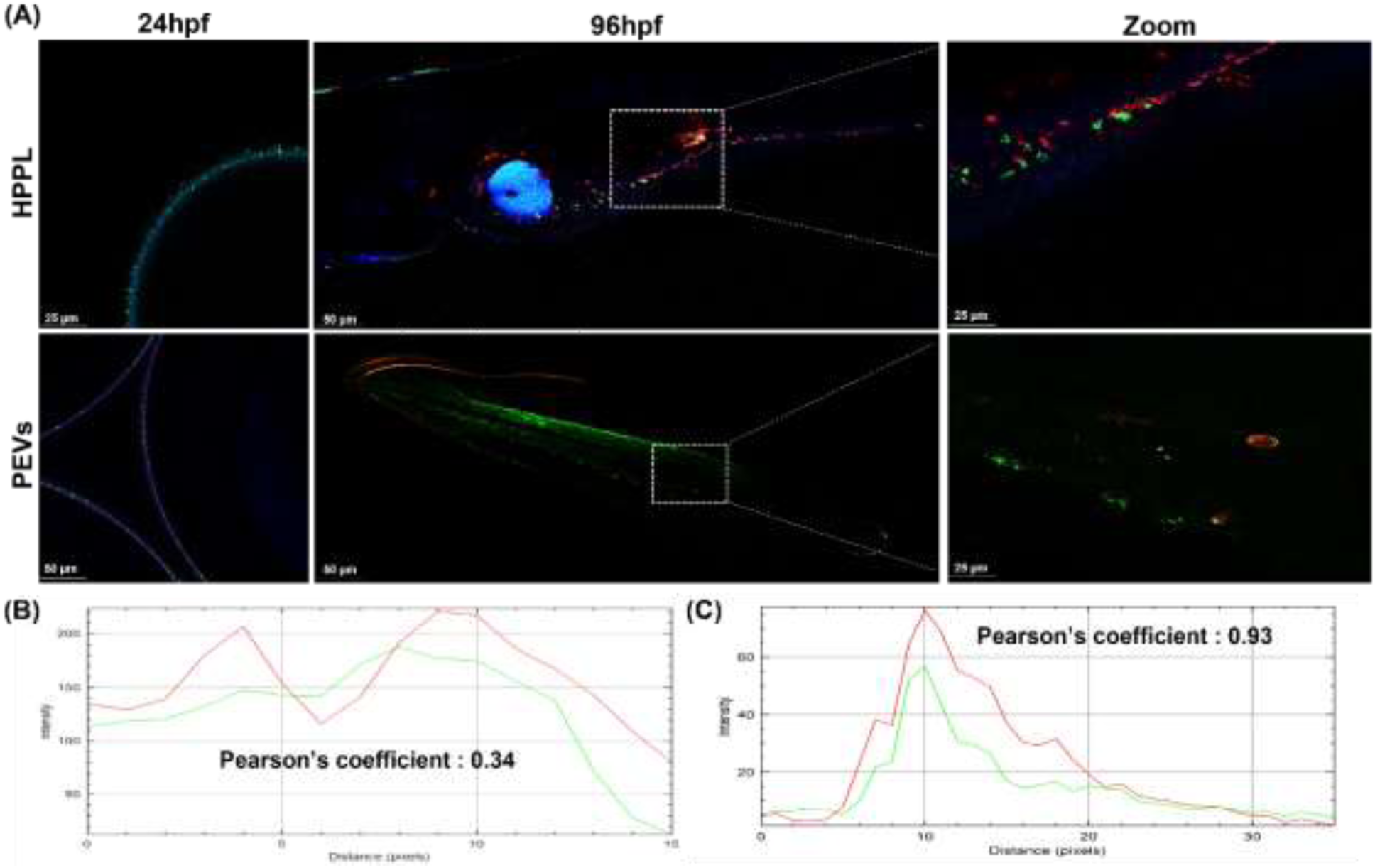
Biodistribution and mitochondrial co-localization of HPPL and PEVs in ZF embryos. (A) Biodistribution of HPPL and PEVs in mitochondria at 24 hpf and 96 hpf, scale bar: 25 μm, 10-fold magnification (B) Quantified Pearson’s coefficient for HPPL colocalization with mitochondria at 96 hpf (C) Quantified Pearson’s coefficient for PEVs colocalization with mitochondria at 96 hpf.

### 3.11. HPPL and PEVs reduce oxidative stress, restore ATP production, and improve mitochondrial morphology in zebrafish embryos

ROS levels detected using DCFDA were markedly elevated in rotenone-treated embryos, indicating cellular oxidative stress. Pre-treatment with HPPL and PEVs reduced ROS signals, suggesting potent antioxidative effects and preservation of redox homeostasis (Figure 12A & B). Next, to further assess mitochondrial function, mitochondria were isolated from ZF embryos after 96 hpf following rotenone exposure. Rotenone treatment caused a pronounced decline by 74.06 % in mitochondrial ATP levels compared to the controls, reflecting severe mitochondrial dysfunction and energy deficit. ZF embryos pretreated with HPPL and PEVs exhibited higher ATP levels in isolated mitochondria compared to the rotenone-only group by 75.38 % and 181.59 %, respectively. This restoration of ATP production suggests that HPPL and PEVs mitigate rotenone-induced mitochondrial damage (Figure 12C). TEM analysis of ZF embryos brain tissue at 96 hpf provided high-resolution visualization of mitochondrial ultrastructure. Embryos exposed to rotenone exhibited pronounced mitochondrial abnormalities, including swelling, disrupted cristae, and outer membrane rupture, hallmarks of severe mitochondrial damage. In contrast, embryos pretreated with HPPL and PEVs showed mitochondria with intact outer membranes and well-preserved cristae, indicating that both treatments effectively protected mitochondrial integrity (Figure 12D). Specifically, rotenone-exposed ZF embryo exhibited an increase in mitochondrial area (averaged 0.70 ± 0.24) as compared to untreated counterparts (0.29 ± 0.079) (Figure 12E). HPPL reduced the mean area by 13% and PEVs treatment significantly reversed by 50% lower as compared to rotenone alone, suggesting better protection against mitochondrial swelling or damage. This enlargement reflects a significant rise in the mitochondrial swelling index, consistent with structural damage. Treatment with HPPL reduced the mean mitochondrial area, while PEVs treatment reversed the effect more robustly, leading to a lower area compared to rotenone alone, suggesting superior protection against mitochondrial swelling (Figure 12F). In addition, cristae score analysis revealed pronounced cristae disruption and rarefaction in rotenone-exposed embryos, whereas HPPL afforded partial preservation. Notably, PEVs treatment substantially restored cristae integrity, yielding significantly higher cristae scores relative to rotenone-treated embryos, supporting their protective role against mitochondrial dysfunction (Figure 12G).

**Figure 12.**
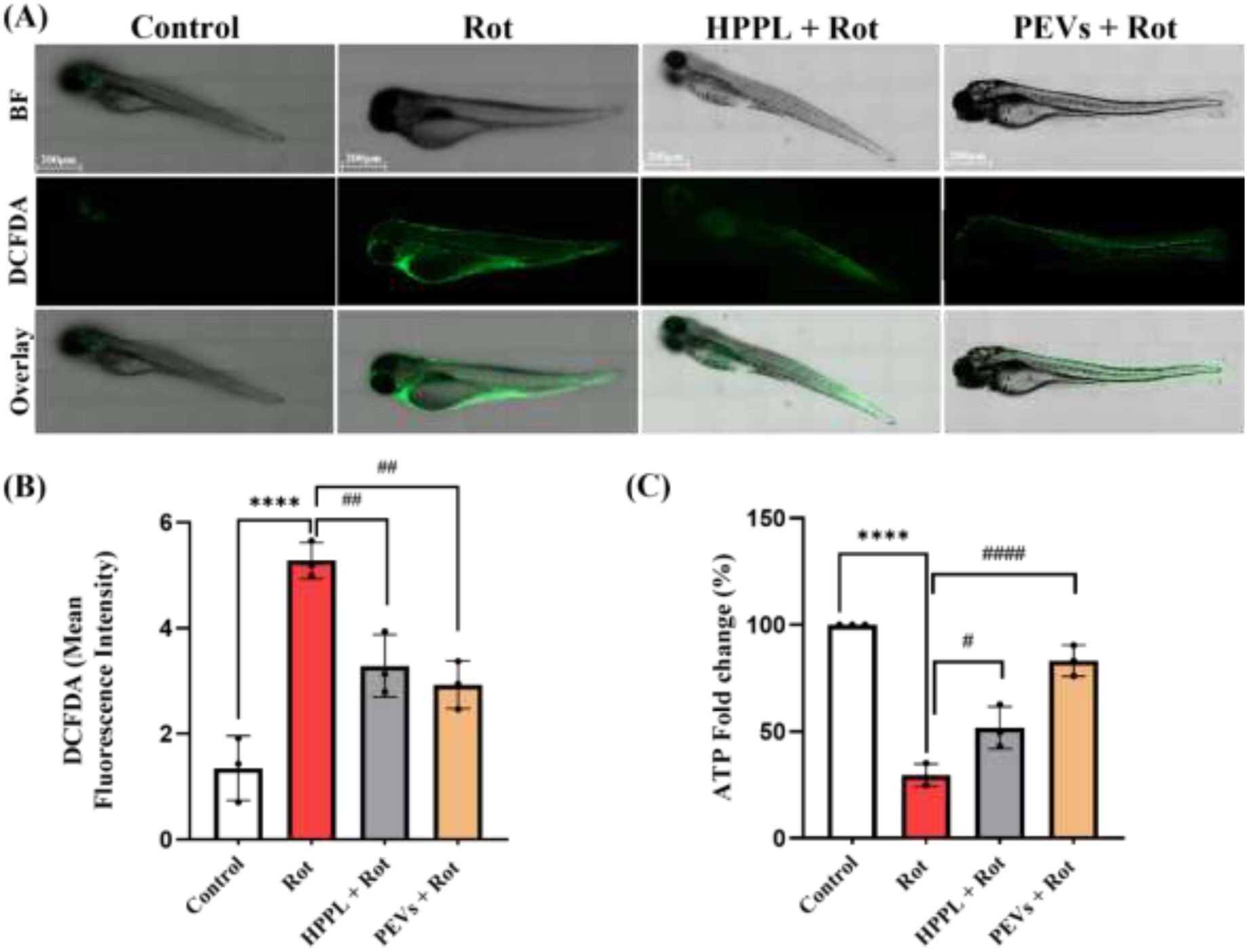
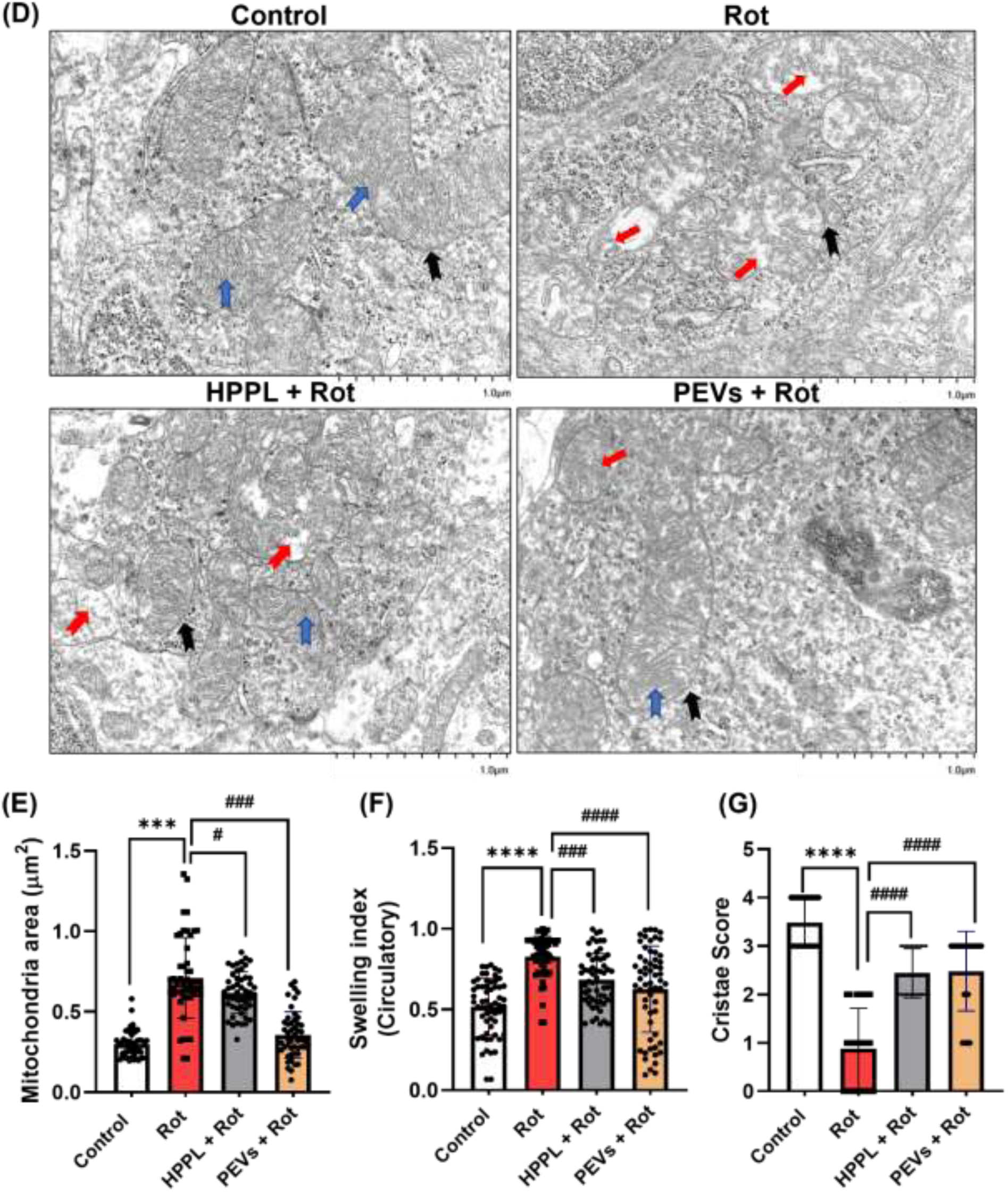
Effect of HPPL and PEVs on oxidative stress, mitochondrial bioenergetics and structure in ZF embryos. (A) ROS generation determined by DCFDA staining, representative images captured at 10-fold magnification, (n = 3) scale bar: 200 μm. (B) Quantification of DCFDA mean fluorescence intensity. (C) mitochondrial ATP production (n = 3). (D) changes in ZF embryo brain tissue mitochondria and cristae observed under TEM, representative images captured at 15-fold magnification, scale bar: 1 μm. Blue arrows represent organized dense mitochondrial cristae, red arrows show collapsed mitochondrial matrix with abnormal cristae, black arrow defines mitochondrial membrane integrity, yellow arrow shows damaged mitochondrial membranes. (E) Quantification of key mitochondrial characteristics included a mitochondrial area (um^2^) (F) Swelling index determined by circulatory which measures mitochondrial shape (< 0.6 elongated/tubular and 1 round/bloated) (G) Measurement of the observed cristae quality. Each dot represents one mitochondrion, with a total number of 50 mitochondria in each condition from at least three biological replicates. Values are expressed as mean ± SD using one-way ANOVA comparisons tests; *P < 0.05, ***P < 0.001, ****P < 0.0001 vs control; ^#^P< 0.05, ^##^P< 0.01, ^###^P< 0.001, ^####^P< 0.0001 vs rotenone group.

## 4. Discussion

Platelet-derived biomaterials are receiving increasing attention as regenerative and neuroprotective agents because of their rich and multimodal bioactive cargo [1, 33]. These materials extend the clinical relevance of platelets far beyond hemostasis by providing trophic, immunomodulatory, antioxidant, and tissue-repairing signals that can be harnessed in cell-free therapeutic formulations [34]. In the present study, we investigated two platelet concentrate-derived biomaterial formulations: HPPL, a soluble lysate generated by controlled platelet disruption and heat treatment [15], and PEVs, a vesicular fraction isolated from platelet concentrate supernatants [9, 35]. Because both are generated from clinical-grade platelet concentrates under defined processing conditions, they offer a clinically relevant, pathogen-screened, and scalable source of biomaterials for translational applications in regenerative medicine and neuroprotection [1, 36–38].

These two biomaterial formats differ structurally and functionally. HPPL provides a soluble pool of platelet-derived trophic and antioxidant factors released after freeze-thaw disruption and subsequent depletion of thrombogenic activity by heat treatment [15]. In contrast, PEVs represent a nanoscale vesicular formulation that preserves membrane architecture and enables the transport of proteins, lipids, and signaling molecules within a lipid bilayer [9, 35]. This distinction is important because it suggests complementary modes of action: HPPL may act as an immediately bioavailable soluble trophic matrix, whereas PEVs may support more selective uptake, trafficking, and intracellular delivery of bioactive cargo. This conceptual distinction strengthens the biomaterial relevance of the study, as it compares two cell-free therapeutic formats derived from the same clinical source material.

The present physicochemical characterization supports the validity of this platform approach. The isolated PEVs displayed the expected vesicular morphology, size distribution, and EV-associated markers, consistent with prior reports and with MISEV-oriented characterization standards [9, 35, 39]. Their origin from pooled clinical-grade platelet concentrates also supports batch consistency and translational manufacturability. In parallel, proteomic profiling of HPPL and PEVs showed that both biomaterials carry a coordinated set of proteins linked to mitochondrial energy metabolism, redox regulation, organelle dynamics, and quality control. The identification of enzymes related to the TCA cycle, OXPHOS, antioxidant defense, and mitochondrial remodeling suggests that platelet-derived biomaterials are not merely trophic supplements, but complex biological matrices capable of modulating several interrelated determinants of mitochondrial homeostasis. In this respect, they behave as multimodal mitochondrial-protective biomaterials rather than single-factor interventions.

A notable translational strength of these platelet-derived biomaterials lies in their clinical source. Clinical-grade PCs, whether obtained by apheresis or by whole-blood processing, are already produced at scale in licensed blood establishments under controlled quality systems [1, 36, 40]. This makes them particularly attractive as starting materials for biomaterial development. In contrast to highly engineered synthetic formulations or cell-based products that require specialized manufacturing, HPPL and PEVs can in principle be prepared from a globally available blood-bank resource. This feature is especially relevant for scalable access, local production, and adaptation to settings with limited advanced manufacturing capacity [17]. Thus, the present study does not only explore biological function, but also supports a practical translational model in which clinically sourced platelet biomaterials may be developed as accessible cell-free therapeutic platforms.

Our in vitro and in vivo findings demonstrate that both HPPL and PEVs counteract rotenone-induced mitochondrial dysfunction and oxidative stress. Rotenone is a well-established mitochondrial complex I inhibitor and induces a robust pattern of mitochondrial injury characterized by impaired OXPHOS, ATP depletion, ROS accumulation, membrane depolarization, and altered ultrastructure [28, 41]. Consistent with this mechanism, rotenone exposure in N2A and SH-SY5Y cells caused marked reductions in cell viability, complex I activity, ATP production, mitochondrial membrane potential, and mitochondrial structural integrity. Both HPPL and PEVs substantially attenuated these alterations, indicating that two distinct platelet-derived biomaterials are able to preserve neuronal mitochondrial function under oxidative stress. This protective effect is likely supported by the antioxidant composition of both biomaterials. HPPL [11] and PEVs [9, 35] contain catalase, superoxide dismutases, glutathione-related enzymes, and other redox-regulatory proteins that are directly relevant to neuronal resilience. Their strong radical-scavenging activity, reflected by ORAC measurements and by restoration of GPx activity in stressed cells, is consistent with the concept that platelet biomaterials reinforce both direct antioxidant buffering and endogenous redox defense systems. This interpretation is further supported by previous *in vivo* work showing that HPPL enhances antioxidant pathways while suppressing NADPH oxidase-dependent oxidative injury after traumatic brain injury [11]. Taken together, these data indicate that platelet-derived biomaterials can act as biologically active redox modulators, a feature that is highly relevant for biotherapy-based neuroprotection.

Beyond antioxidant effects, the current study shows that platelet-derived biomaterials influence mitochondrial bioenergetics and structural remodeling. Both HPPL and PEVs restored ATP production and preserved mitochondrial membrane potential, while HPPL additionally showed a stronger effect on maximal respiration. These findings suggest that the protective activity of platelet biomaterials is not limited to ROS scavenging, but extends to preservation of functional oxidative metabolism. Proteomic analyses further support this interpretation by revealing restoration of proteins linked to oxidative phosphorylation, TCA cycle-related pathways, antioxidant defense, and mitochondrial dynamics in rotenone-stressed cells following pretreatment. Thus, and interestingly, the biological activity of HPPL and PEVs appears to result from coordinated actions on multiple mitochondrial stress-response pathways rather than from a single dominant mechanism.

The two platelet-derived biomaterial formats, however, were not identical in their effects. HPPL showed stronger effects on mitochondrial respiration and redox recovery, whereas PEVs more prominently influenced some aspects of mitochondrial remodeling and in vivo protection. This divergence is biologically plausible and biomaterial-relevant. HPPL is rich in soluble growth factors, chemokines, and antioxidants, including BDNF, VEGF, PDGF, PF4/CXCL4, CCL5, TGF-β, EGF, and HGF [8]. Many of these factors promote neuronal survival, differentiation, and repair [37, 42], and some, such as BDNF and IGF-1, are linked to mitochondrial regulation [42, 43]. PEVs, in contrast, provide a vesicle-associated cargo that may enable different uptake dynamics and intracellular bioavailability [44]. The comparison between HPPL and PEVs therefore highlights an important biomaterials principle: materials derived from the same biological source may display distinct functional profiles depending on whether their cargo is delivered as a soluble matrix or within membrane-enclosed vesicles. This observation echoes our recent findings where we observe distinct pro-neurogenic activities of HPPL and of another formulations of PEVs [16]. It also supports that the emerging conceot of the feasibility of engineering specific platelet derived products and secretome preparations for precision neuromedicine [3].

The uptake studies are consistent with this interpretation. Both HPPL and PEVs were internalized by neuronal cells, but PEVs showed greater spatial proximity to mitochondria. Although this does not prove direct mitochondrial targeting, it supports the possibility that vesicular biomaterials may engage distinct intracellular trafficking pathways compared with soluble platelet lysates. At the same time, the limited mitochondrial colocalization observed in neuronal cells suggests that much of the protective activity of PEVs may still be mediated through cytosolic or signaling-based mechanisms rather than direct organelle docking. The contrast with other reports showing mitochondrial localization of extrusion-derived PEVs in different cell types [45] suggests that intracellular trafficking may depend on the EV preparation method, recipient-cell identity, and stress context. This remains an important question for future biomaterial optimization.

Another key finding is that platelet-derived biomaterials modulate mitochondrial quality-control pathways. Both HPPL and PEVs restored the expression of PGC-1α, MFN1, and DRP1, indicating improved mitochondrial biogenesis and normalization of fusion-fission balance. Because mitochondrial dynamics are tightly linked to respiration, ROS handling, and stress adaptation [46–49], these findings are highly relevant to the development of biomaterials for mitochondrial rescue. The restoration of MFN1 and the improvement in mitochondrial ultrastructure indicate preservation of network integrity, while normalization of DRP1 suggests that the benefit is not limited to enhanced fusion but involves broader rebalancing of mitochondrial remodeling. In this respect, HPPL and PEVs behave as dynamic mitochondrial-supportive biomaterials capable of stabilizing structure-function relationships under pathological stress.

The in vivo zebrafish data further strengthen this conclusion. Zebrafish embryos provide a useful and accessible system for evaluating mitochondrial injury, developmental toxicity, oxidative stress, and early brain effects in a transparent and experimentally tractable vertebrate model [50, 51]. In this study, both platelet biomaterials improved embryo survival, hatching, oxidative stress markers, and mitochondrial ultrastructure after rotenone exposure, with PEVs showing particularly strong efficacy. These data indicate that the protective effects of HPPL and PEVs are not restricted to cultured neurons but extend to a whole-organism injury context, also consistent with preclinical studies conducted in rodents [3, 12]. The observed protection in embryonic brain mitochondria is particularly relevant, as it supports the idea that platelet-derived biomaterials retain activity in complex biological environments.

The stronger performance of PEVs in some zebrafish endpoints may reflect vesicle-specific biodistribution and trafficking properties. Fluorescent imaging suggested broader tissue uptake and stronger mitochondrial colocalization of PEVs in embryos, which may enhance delivery efficiency in vivo. This is consistent with previous reports showing that platelet-derived EVs display context-dependent homing and biodistribution patterns that vary with platelet activation and vesicle composition [52]. Such properties may give vesicular platelet biomaterials an advantage in certain translational settings where tissue penetration, local retention, or intracellular delivery are important. By contrast, the soluble nature of HPPL may favor broader extracellular trophic support. Together, these observations suggest that the two biomaterial formulations may have complementary translational applications rather than being interchangeable, again supporting our recent findings in rodent hippocampal neurogenesis *ex vivo* and *in vivo* [53].

From a therapeutic perspective, this complementarity is one of the most interesting outcomes of the study. HPPL appears particularly effective for soluble trophic support, antioxidant buffering, and respiratory recovery, whereas PEVs may provide additional benefits linked to vesicle-mediated trafficking, mitochondrial remodeling, and in vivo resilience. This raises the possibility that combined or sequential use of platelet-derived soluble and vesicular biomaterial formats could enhance neuroprotective efficacy. Such a strategy would be fully consistent with current biotherapies thinking, where multifunctional or complementary platforms are increasingly favored over single-component interventions [3, 54–56].

The study also has limitations. Rotenone is a robust model of mitochondrial complex I inhibition, but it does not fully capture the disease-specific complexity of Parkinson’s disease or other neurodegenerative disorders. In addition, the present work focused mainly on complex I-related injury and did not systematically examine effects on other respiratory complexes. While the in vivo zebrafish data are informative, more disease-specific mammalian models and longer-term functional outcomes will be needed to fully define translational potential. Further work should also clarify cargo-function relationships, determine how bioprocessing influences biomaterial activity, and identify which soluble or vesicular components most strongly drive mitochondrial rescue. These questions are important not only biologically but also for future biomaterial optimization, manufacturing control, and regulatory translation.

In summary, the present study supports the concept that HPPL and PEVs are two clinically sourced platelet-derived biomaterial formulations with complementary mitochondrial-protective concept summarized in supplementary figure 7. Their shared origin from clinical-grade platelet concentrates, combined with their distinct cargo architectures and functional profiles, makes them attractive candidates for scalable cell-free neuroprotective development. By integrating antioxidant, metabolic, and mitochondrial quality-control effects, these platelet-derived biomaterials provide a strong rationale for further translational exploration in neurodegenerative disorders.

<u>/</u>

## 4. Conclusion

In conclusion, this study highlights the potential of both HPPL and PEVs as promising platelet-derived biomaterials for mitigating mitochondrial dysfunction associated with neurodegeneration. Both biomaterials reduced oxidative stress, preserved mitochondrial function, and improved cellular bioenergetics in rotenone-exposed neuronal cells and zebrafish embryos. Their origin from clinical-grade platelet concentrates provides a scalable and pathogen-screened resource for <u>the development of accessible cell-free biotherapies.</u>

<u>*</u>

## Abbreviations

AM: Axis malformation
ATP: Adenosine triphosphate
AD: Alzheimer&#x2019;s disease
ALS: Amyotrophic lateral sclerosis
BT: Bent tail
CNS: Central Nervous system
Cryo-EM: Cryo-electron microscopy
DLS: Dynamic Light scattering
ETC: electron transport chain
EVs: extracellular vesicles
GSTs: glutathione S-transferase
GPx: glutathione peroxidase
hpf: hour post fertilization
HPPL: heat-treated platelet pellet lysates
HPL: human platelet lysates
MFN-1: Mitofusin-1
MitoQ: MitoQuinone Mesylate
MPTP: mitochondrial permeability transition pore
NADPH: Nicotinamide adenine dinucleotide phosphatase
NTA: Nanoparticle tracking assay
OXPHOS: Oxidative phosphorylation cycle
OCR: oxygen consumption rate
PCs: platelet concentrates
PD: Parkinson’s disease
PE: pericardial edema
PFA: paraformaldehyde
PGC1a: peroxisome proliferator-activated receptor gamma coactivator 1a
PEVs: platelet derived extracellular vesicles
PPL: platelet pellet lysates
ROS: reactive oxygen species
SIRT1: Sirtuin-1
SOD: Superoxide dismutase
TBI: traumatic brain injury
TEM: Transmission electron microscopy
TFAM: mitochondrial transcription factor A
TOMM20: Translocase of Outer Mitochondrial Membrane 20
YE: yolk sac edema axis

## Declarations

### Ethics approval and consent to participate

The Institutional Review Board of Taipei Medical University approved this study (TMU-JIRB N201802052).

### Availability of data and materials

Data analyzed during the current study are available from the corresponding authors upon reasonable request.

### Author contributions

**KG**: Conceptualization, Data Curation, Formal Analysis, Investigation, Methodology, Validation, Writing- Original Draft, Reviewing and Editing. **LD**: Methodology, Data Curation, Formal Analysis, Investigation, **MLC**: Formal Analysis, Software, Validation, Visualization, Writing-Original Draft, Review and Editing **LTNN:** Methodology, Investigation, Software, Validation. **HYW:** Methodology, Investigation. **AP:** Conceptualization, Supervision, Writing- Reviewing and Editing **DB**: Formal Analysis, Methodology, Writing- Review and Editing. **TB**: Conceptualization, Validation and Supervision, Formal Analysis, Funding Acquisition, Project Administration, Resources, Writing- original draft, review and editing.

## Acknowledgements

We thank the Core Imaging Facility, TMU, for their help with the sample sectioning for TEM analysis, the zebrafish Core Lab Facility, TMU, for providing the zebrafish embryos, the Consortia of Key Technologies and Instrumentation Center at National Taiwan University for providing mass spectrometry technical services for proteomics analysis, and the Taipei Blood Center (Taiwan Blood Service Foundation, Guandu, Taiwan) for supplying the PCs for this study.

