## Supplementary figure 1-7 for "Clinical-grade platelet-derived biomaterials mitigate mitochondrial dysfunction in preclinical models of neuronal injury"

*** Corresponding Authors:**

orcid.org/0000-0002-0507-9243

1. **Platelet surface markers expressing CD62P, CD42A and CD41 in four different batches of isolated PEVs.**

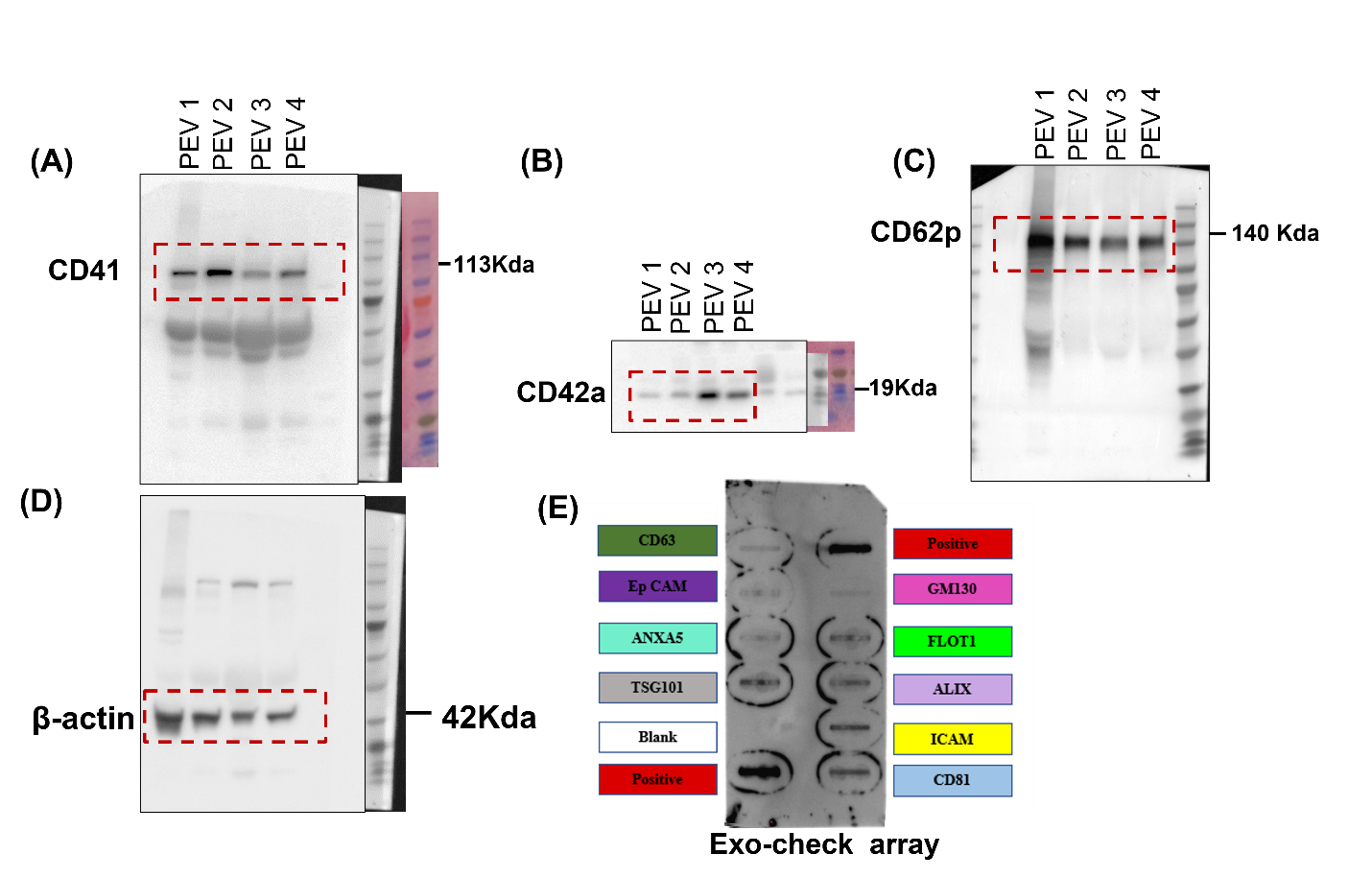

**Supplementary figure 1:** Uncropped images of different batches of PEVs western blot for platelet surface markers **(A)** CD41, **(B)** CD42a, **(C)** CD62p **(D)** β-actin protein levels; βactin was used as loading control **(E)** Original image of the human exosome antibody array after incubation with pooled sample of PEVs, showing EV markers and controls.

2. **Preliminary tests for dose standardization for rotenone, HPPL and PEVs**

To establish appropriate working concentrations for the main experiments, preliminary dose-response assays were conducted in N2A and SH-SY5Y neuronal cell lines. Both cells were exposed to increasing concentrations of rotenone (0.5–10 µM) for 24 hr. A dose-dependent reduction in cell viability was observed with rotenone, with cytotoxicity evident at concentrations ≥ 3 µM. Based on these results, 5 µM rotenone was selected for subsequent experiments. Next, the effects of platelet-derived biomaterials were evaluated: HPPL and PEVs were applied to N2A cells and SH-SY5Y cells at escalating doses (1%–5% *v/v*) for 24 hr. Both cell lines exhibited a dose-tolerant profile with no significant reduction in viability up to 5% (*v/v*) for either HPPL or PEVs. Higher concentrations (5%v/v) were selected as the optimal non-toxic and biologically active dose for subsequent in vitro studies.

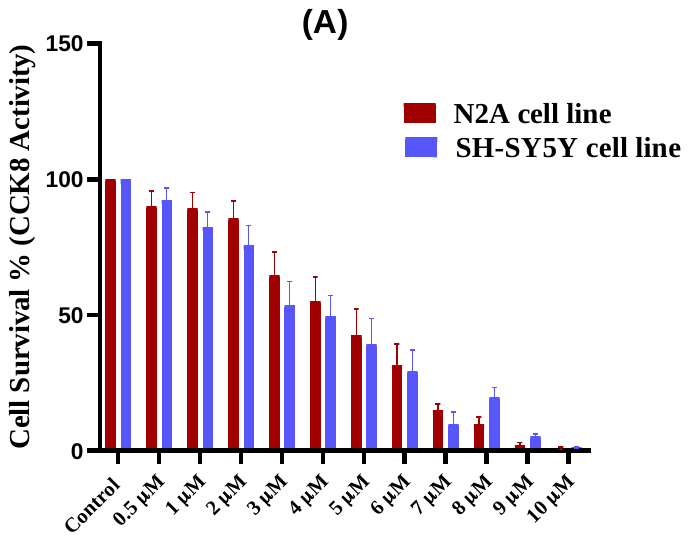

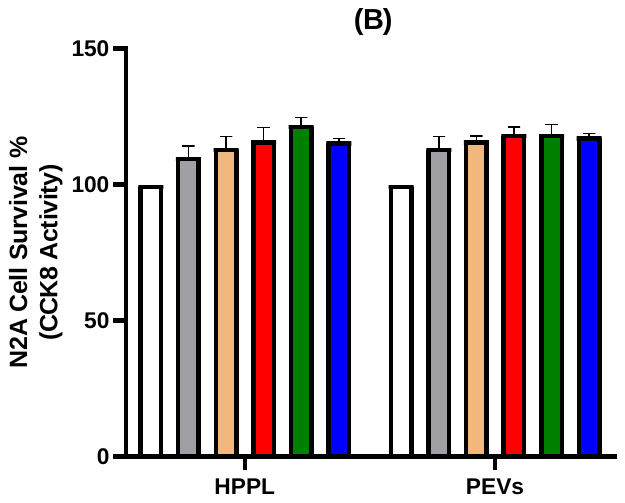

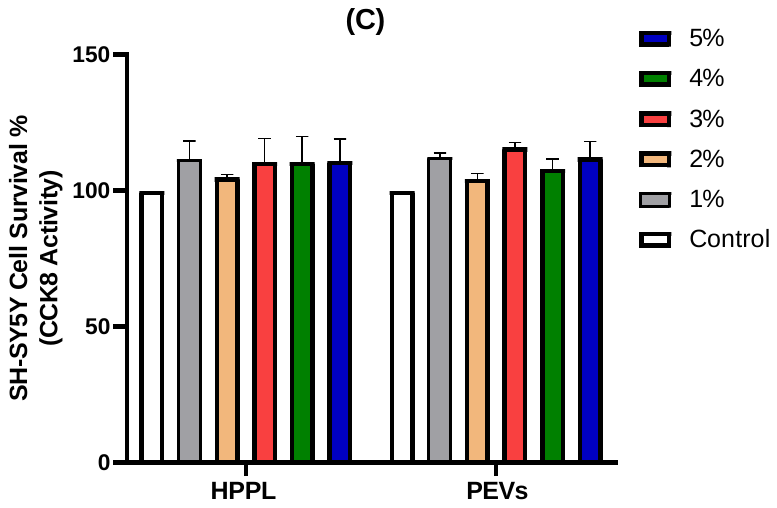

**Supplementary Figure 2:** Pre-liminary testing for dose response study of (A) Rotenone, (B) HPPL and PEVs incubated for 24 hours in N2A cells (D) HPPL and PEVs incubated for 24 hours in SH-SY5Y cells. Results expressed as a mean ± SD from n=6 biological replicates.

3. **Mitochondrial electron transport chain (ETC) complexes and their differential regulation across experimental groups identified by Ingenuity Pathway Analysis (IPA)**
(A) **Rotenone Vs Control**

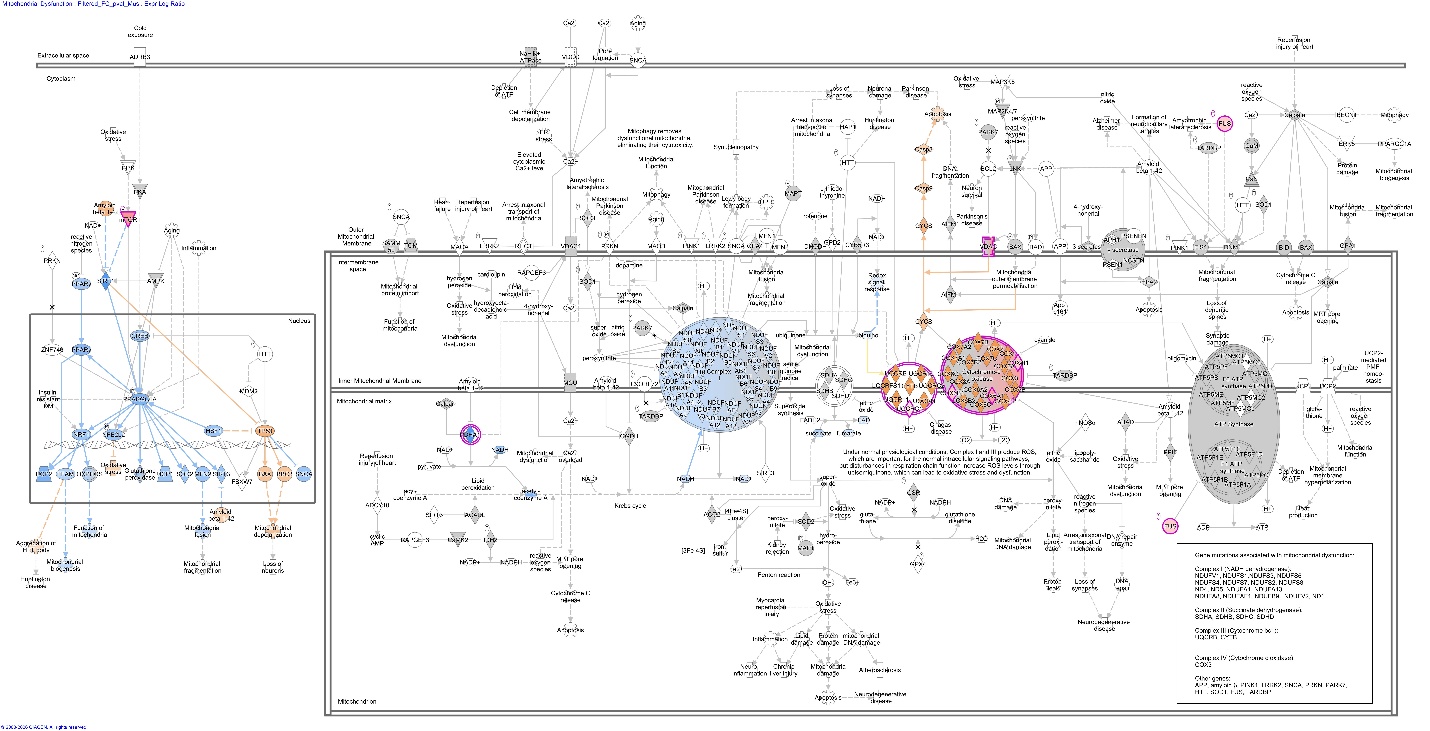

(B) **Rotenone + HPPL vs Rotenone**

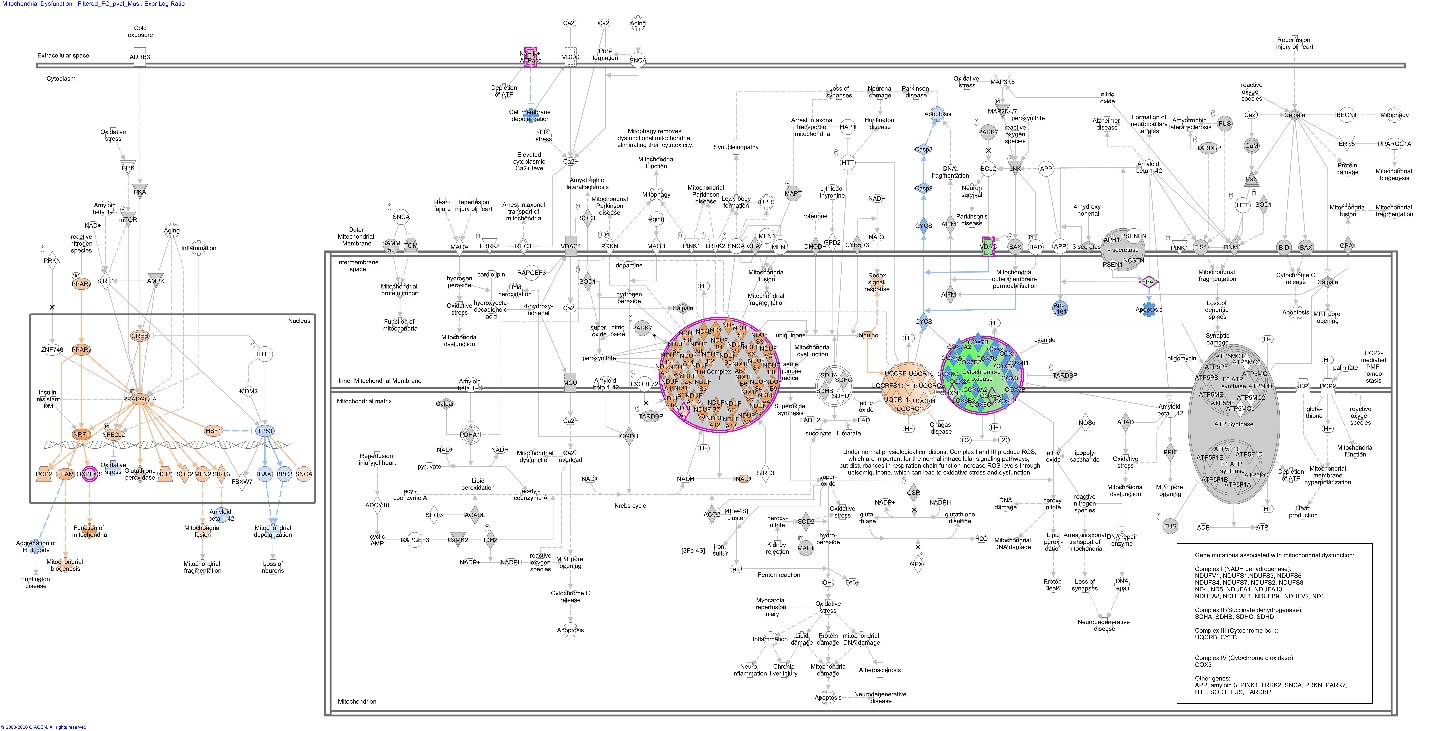

(C) **Rotenone + PEVs vs Rotenone**

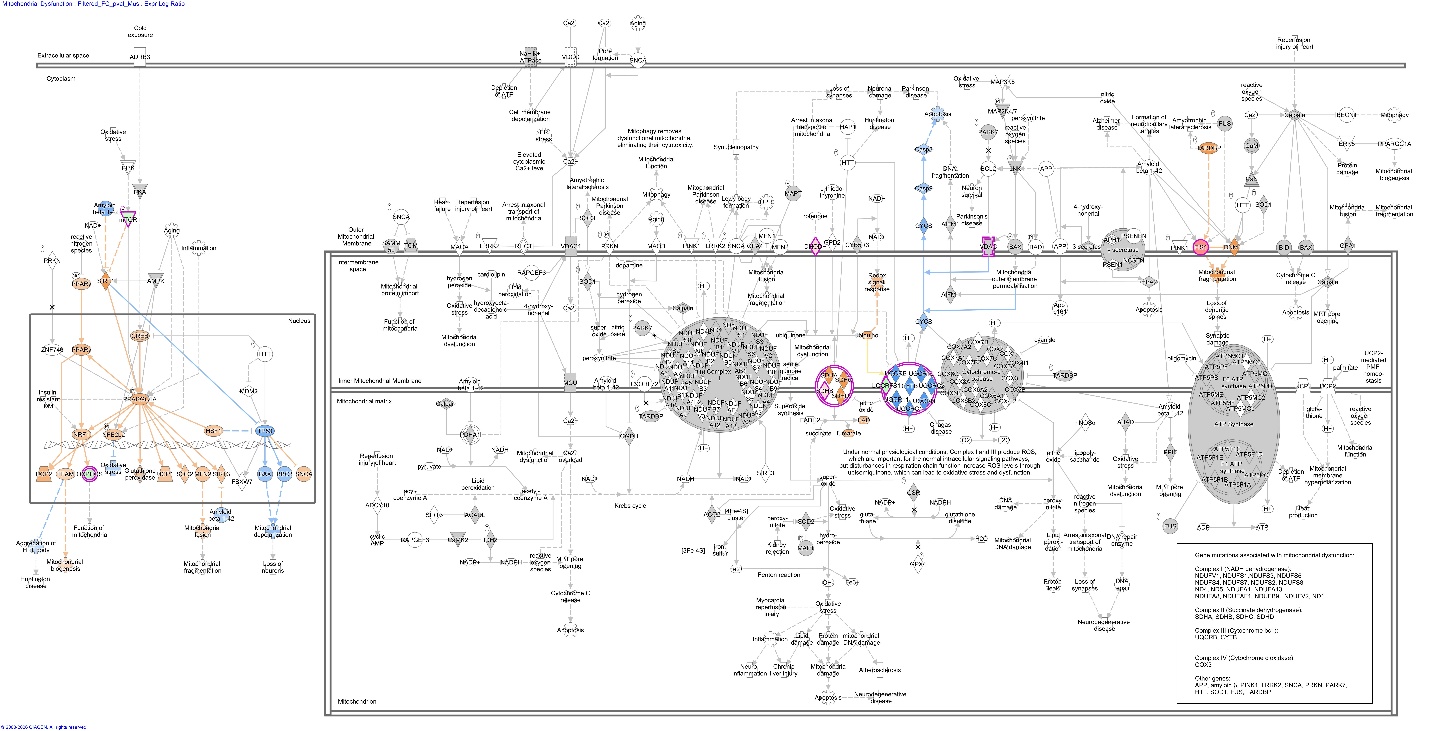

**Supplementary Figure 3:** **Representation of mitochondrial electron transport chain (ETC) complexes and their differential regulation across experimental groups identified by Ingenuity Pathway Analysis (IPA)**
Schematic representation of the mitochondrial **electron transport chain (ETC)** including **Complex I–V,** which are responsible for oxidative phosphorylation and ATP production. Differential protein expression associated with mitochondrial pathways was analyzed using Ingenuity Pathway Analysis. The diagram illustrates changes in ETC-associated proteins across experimental comparisons**: (A) Rotenone vs Control, (B) Rotenone + HPPL vs Control,** and **(C) Rotenone + PEVs vs Control**. Proteins are color-coded according to their expression levels, where **red indicates upregulation, blue indicates downregulation,** and **grey indicates no significant change.** The analysis highlights alterations in mitochondrial bioenergetics induced by rotenone treatment and the modulatory effects of HPPL and PEVs on mitochondrial function.

4. **Preliminary assessments in Zebrafish Embryo:**

**Dose validation for rotenone:** The secondary objective of this study was to develop the ZF as a platform for testing the in vivo biological effects of rotenone, we therefore sought to assess the feasibility of using ZF as a model for mitochondrial dysfunction. Following our in vitro studies, we examined ZFET test for different exposures of HPPL, PEVs and rotenone concentrations. Based on these assessments, we selected 200 nM rotenone as optimal concentrations for subsequent experiments. This approach allowed us to effectively model mitochondrial impairment in a living ZF embryos

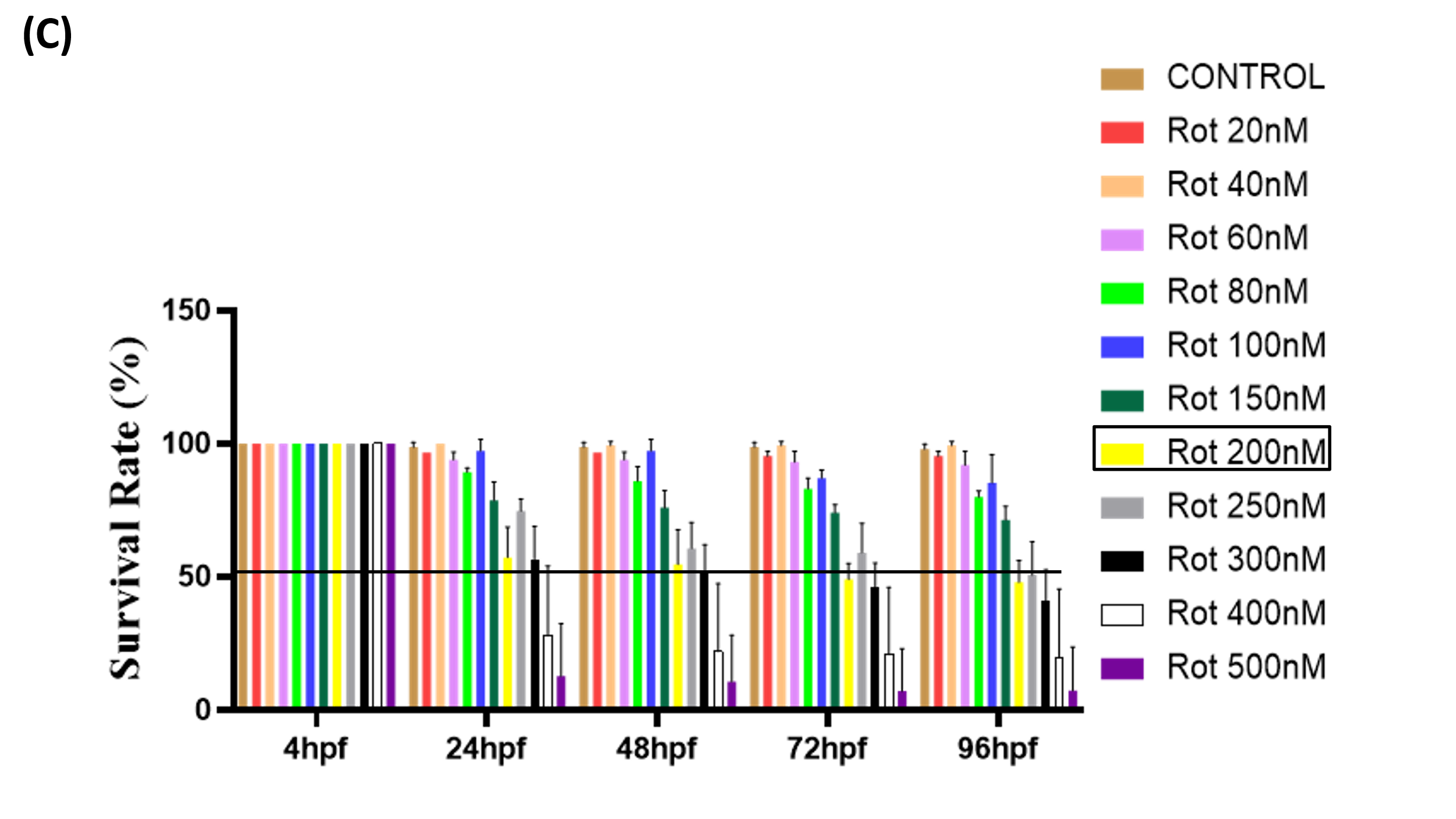

**Supplementary Figure 4:** Survival percentage of ZF embryo at 4, 24, 48, 72, 96 hpf at different concentrations of rotenone (20 nM-500 nM).

5. **Optimization of HPPL and PEVs Dosage Based on Protein and Particle Load in Zebrafish Embryos**

To ensure biologically relevant and non-toxic exposure conditions, PEVs concentrations for zebrafish embryo experiments were optimized based on both protein content and particle count. Preliminary findings indicated that protein doses exceeding 30 µg per mL in embryo medium led to adverse developmental effects, establishing an upper threshold for therapeutic protein delivery. Accordingly, PEVs (20 μg/mL) was optimized, which was within the safe and effective range for zebrafish embryo exposure. Additionally, nanoparticle tracking analysis (NTA) revealed that undiluted PEVs stocks often contained 10¹¹–10¹² particles/mL, and using high-particle suspensions directly in E3 medium will be potentially disruptive to ZF embryo development. By coordinating both protein and particle parameters, our final dosing strategy balanced therapeutic efficacy with biocompatibility, enabling consistent, reproducible in vivo evaluation of PEVs effects in zebrafish embryos.**v**Similar to PEVs, the dosing of HPPL for zebrafish embryo treatment was optimized to avoid protein overload while maintaining therapeutic efficacy. To ensure that the protein exposure remained below the empirically defined 30 µg/mL threshold, HPPL was diluted, resulting in a final protein concentration of approximately 20-25 µg/mL. This dilution strategy allowed effective delivery of bioactive factors from HPPL while minimizing developmental toxicity in zebrafish embryos. Unlike PEVs, particle concentration was not a limiting factor for HPPL application. Therefore, protein-based dosing served as the primary guide for safe and standardized HPPL administration in the zebrafish model.

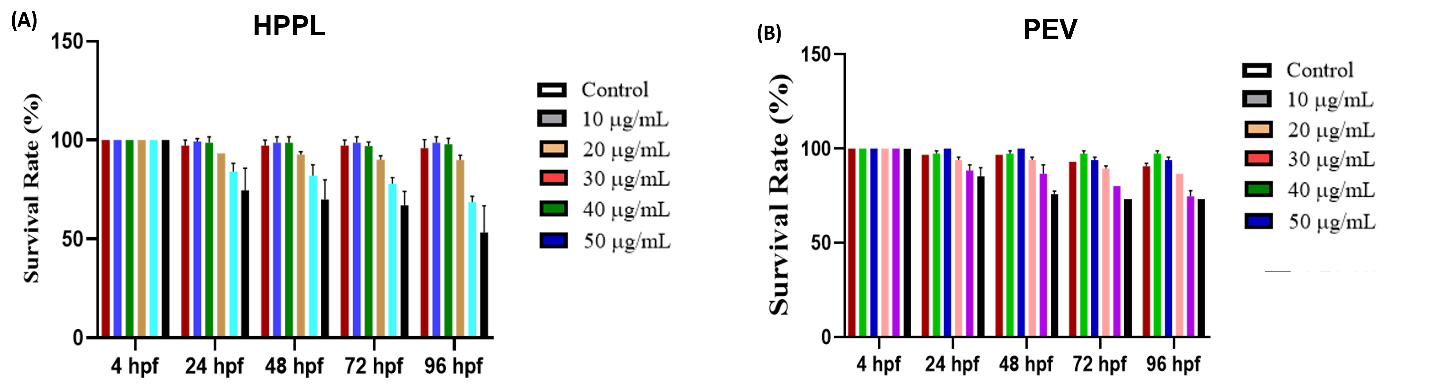

**Supplementary Figure 5:** Survival percentage of ZF embryo at 4, 24, 48, 72, 96 hpf at different concentrations of **(A)** HPPL and **(B)** PEVs.

6. **Individual quantification of developmental abnormalities in zebrafish embryos**

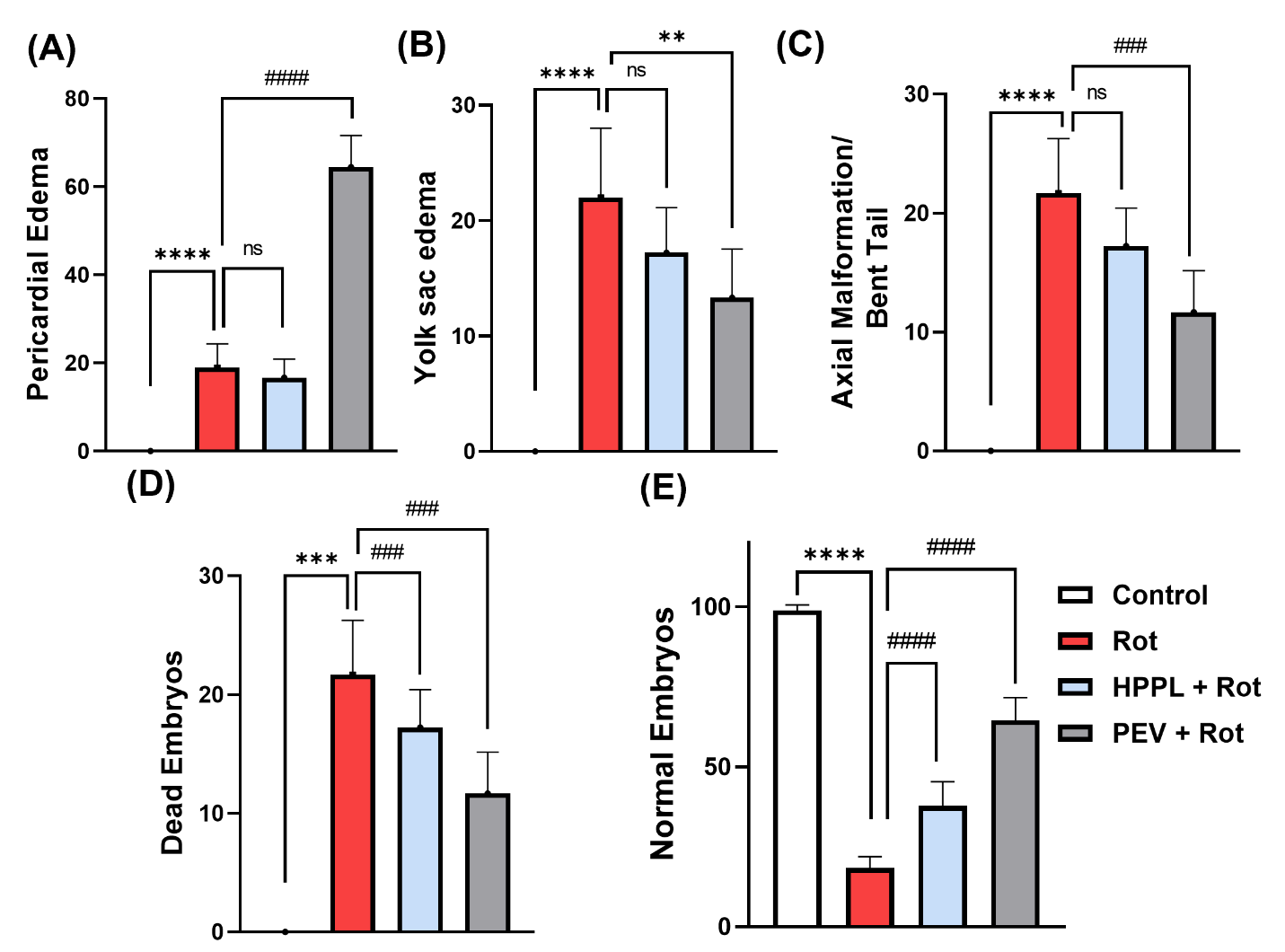

**Supplementary Figure 6:** **Quantification of developmental abnormalities including (A)** Pericardial edema**, (B)** Yolk sac edema**, (C) Axial Malformation,** Bent tail **(D) Dead Embryos(E)** Normal Embryos (% affected embryos per group), compiled from six independent experiments. Values are expressed as mean ± SD (n = 6); ****P < 0.0001 vs control; ^###^P < 0.001 vs rotenone treated embryos

7. **Detailed schematic representation of rational, plan of work and summary of the current study**

Platelets were selected as a source because they are natural reservoirs of bioactive molecules, including antioxidants, trophic factors, and extracellular vesicles that support mitochondrial integrity. Prepared under clinical-grade conditions, platelet concentrates provide a safe, standardized, and readily available material for therapeutic development. Unlike cell therapies, platelet-derived lysates and EVs are acellular, minimizing immunogenicity while delivering high levels of GPx, SOD, catalase, and neurotrophic factors. These components counteract oxidative stress, preserve mitochondrial function, and promote neuronal survival. Their established safety in transfusion and regenerative medicine further underscores their translational relevance. Together, this positions HPPL and PEVs as promising, scalable cell-free biotherapies for restoring redox balance and protecting against neurodegeneration. Focused experiments on ETC complexes and mitochondrial dynamics should be undertaken in future to establish specific roles of HPPL and PEVs therapy.

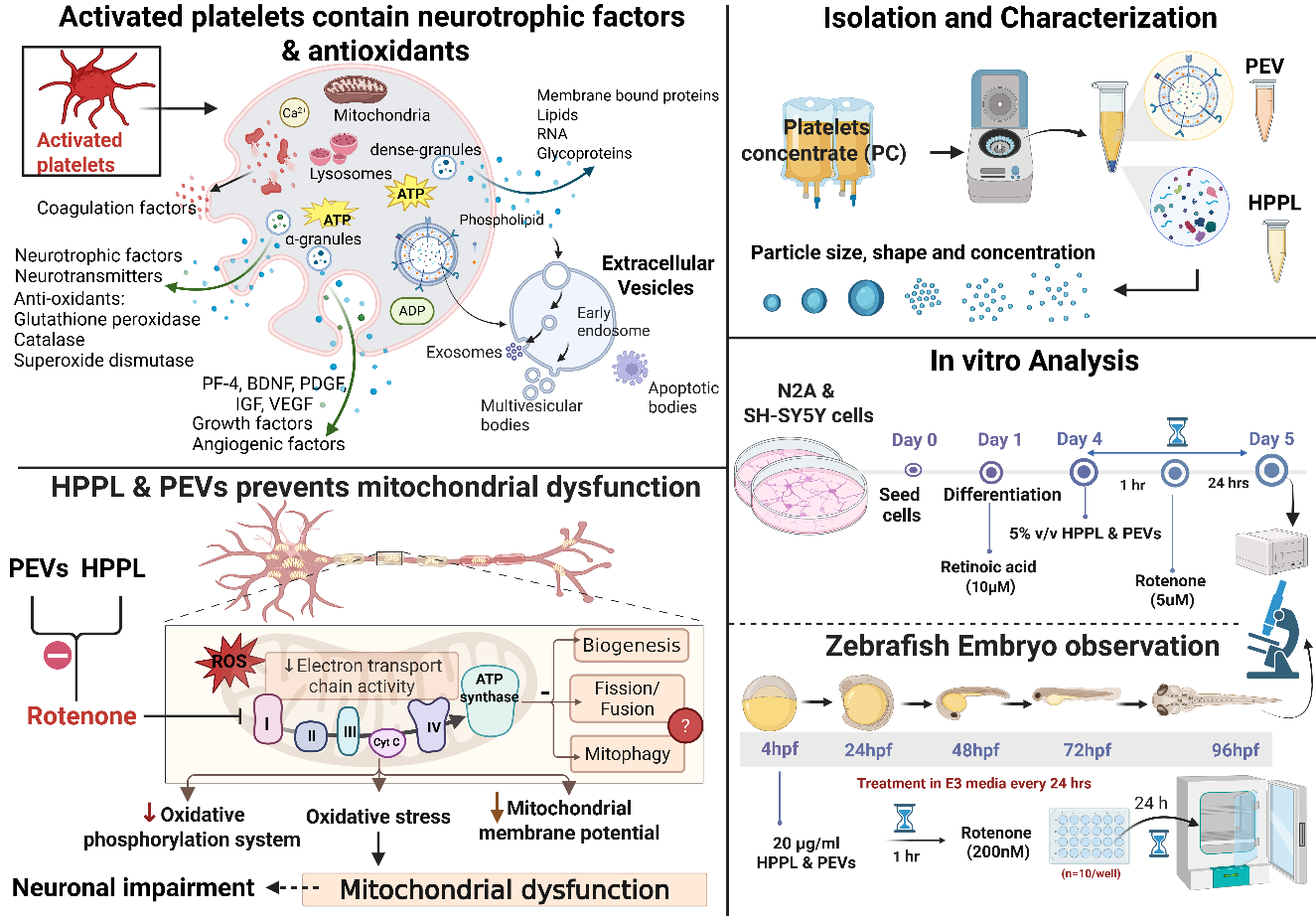

**(A)**

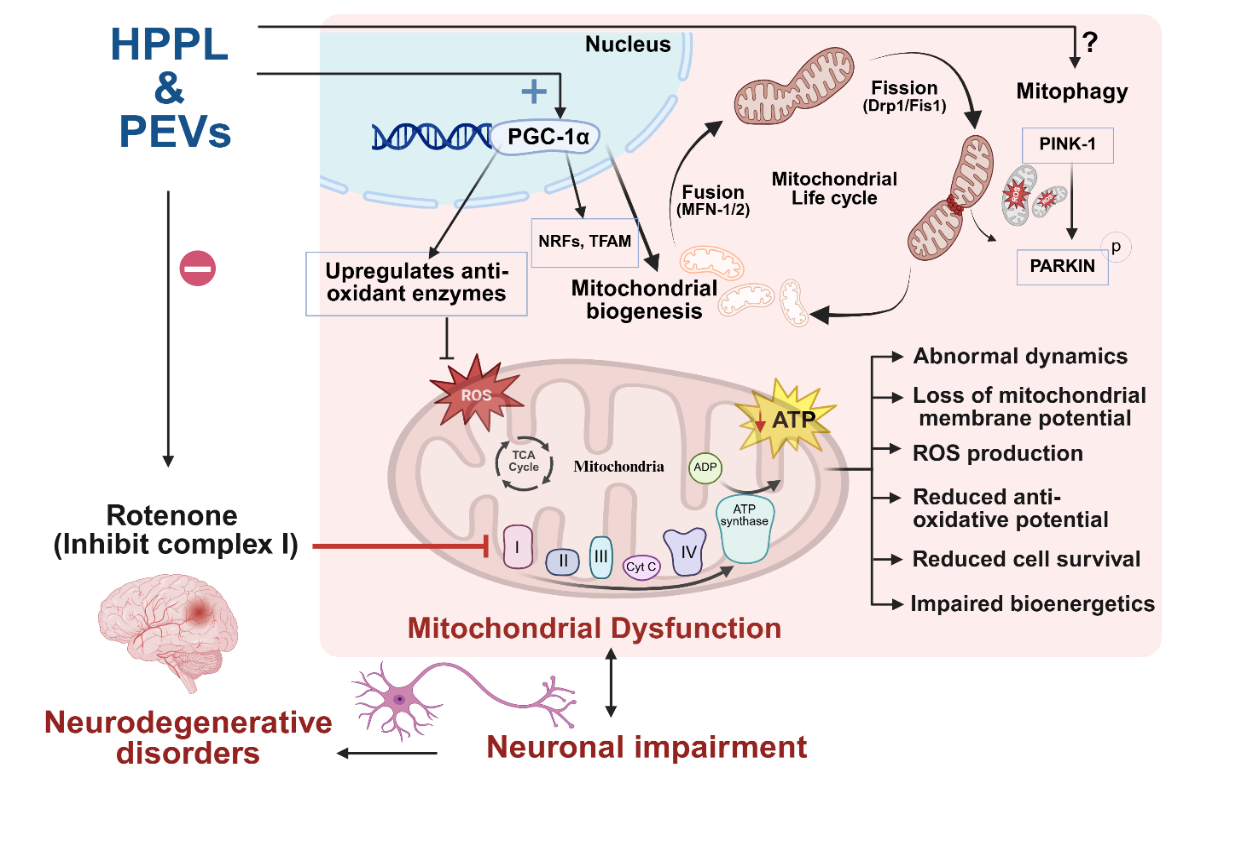

**(B)**

**Supplementary figure 7:** Schematic representation of HPPL and PEVs as neuroprotective agents against rotenone-induced mitochondrial dysfunction. **(A)** Activated platelets release antioxidant molecules and bioactive factors, platelet lysate and EVs are isolated and comprehensively characterized using nanoparticle tracking analysis, transmission electron microscopy, and molecular marker profiling to confirm their physicochemical properties and bioactive content, functional evaluation in vitro and in vivo, endpoints included survival and hatching rates, morphological assessment, ATP levels, and reactive oxygen species (ROS) production. Overall, the schematic illustrates the workflow and highlights the role of PEVs in delivering growth factors to mitigate oxidative stress, restore mitochondrial function, and promote neuroprotection. **(B**) Rotenone inhibits complex I of the ETC, leading to oxidative stress, loss of membrane potential, ROS accumulation, and impaired mitochondrial dynamics, ultimately causing neuronal damage. HPPL and PEVs treatment restore mitochondrial function, reduce oxidative damage, and promote biogenesis and dynamics, highlighting their neuroprotective potential. Created in <https://BioRender.com>.
